# Cytosolic termini of the FurE transporter regulate endocytosis, pH-dependent gating and specificity

**DOI:** 10.1101/588368

**Authors:** Georgia F. Papadaki, George Lambrinidis, Andreas Zamanos, Emmanuel Mikros, George Diallinas

**Affiliations:** Department of Biology, National and Kapodistrian University of Athens, Panepistimioupolis, Athens 15784, Greece.; Department of Pharmacy, National and Kapodistrian University of Athens, Panepistimioupolis, Athens 15771, Greece.

**Keywords:** *Aspergillus nidulans*, fungi, transport, folding, membrane lipids, allosteric

## Abstract

FurE, a member of the NCS1 family, is an *Aspergillus nidulans* transporter specific for uracil, allantoin and uric acid. Recently we showed that C- or N-terminally truncated FurE versions are blocked for endocytosis and, surprisingly, show modified substrate specifities. Bifluorescence complementation assays and genetic analyses supported that the C- and N-termini interact dynamically and through this interaction regulate selective substrate translocation. Here we functionally dissect and delimit distinct motifs crucial for endocytosis, transport activity, substrate specificity and folding, in both cytosolic termini of FurE. Subsequently, we obtain novel genetic and *in silico* evidence supporting that the molecular dynamics of specific N- and C-terminal regions affect allosterically the gating mechanism responsible for substrate selection, via pH-dependent interactions with other internal cytosolic loops and membrane lipids. Our work shows that elongated cytoplasmic termini, acquired through evolution mostly in eukaryotic transporters, provide novel specific functional roles.

## Introduction

Transporters are membrane proteins that mediate the import and export of nutrients, metabolites, signaling molecules or drugs in and out of cells, and are thus essential for cell communication and life. Despite their evolutionary, structural and functional differences all transporters use an alternating-access mechanism where a substrate binding site, in allosteric co-operation with distinct gating domains, alternates between multiple conformations for receiving and delivering specific substrate(s) from one side of the membrane to the other. This basic mechanism, carried out by dynamic movements of the main transmembrane body and assisted by the flexibility of interconnecting hydrophilic loops, exists in different variations, known as the rocker-switch, the rocking-bundle or the elevator sliding mechanisms (Krishnamurthy et al, 2009; Kaback et al, 2011; Drew and Boudker 2016; Kazmier et al, 2017).

One of the best-studied families of transporters is the so-called Nucleobase Cation Symporter 1 (NCS1) family. This is due to a plethora of genetic and biochemical findings concerning fungal members of the family (Krypotou et al, 2012; 2015; Sioupouli et al, 2017), as well as, extensive structural and biophysical data concerning a bacterial homologue, the benzyl-hydantoin/Na^+^ Mhp1 symporter (Weyand et al, 2008; Shimamura et al, 2010; Simmons et al, 2014). NCS1 proteins consist of 12 transmembrane α-helical segments (TMSs) interconnected with rather short loops and cytosolic N- and C-termini. TMSs 1-10 are arranged as a 5-helix intertwined inverted repeat (5HIRT), the so called LeuT-fold, also found in other transporter families involved in neurotransmitter, sugar, amino acid, and drug transport (Shi, 2013; Västermark and Saier, 2014; Drew and Boudker, 2016). The last two TMSs (11 and 12) seem to be crucial for the oligomerization state of some NCS1-like transporters, rather than being involved in the mechanism of transport (Ilgü et al, 2016).

The crystal structures available for Mhp1 correspond to conformational distinct outward-facing open, substrate-occluded and cytoplasm-open topologies (Weyand et al, 2008; Shimamura et al, 2010; Simmons et al, 2014), strongly supporting an alternating access rocking-bundle transport mechanism (Forrest and Rudnick, 2009; Adelman et al, 2011; Simmons et al, 2014; Kazmier et al, 2014). In all cases, TMSs 1, 2, 6, and 7 form a four helix bundle, while TMSs 3, 4, 8, and 9 form a motif that resembles a hash sign. The substrate and Na^+^ binding sites are found between the hash and bundle motifs and involve residues in TMSs 1 and 6, at a position where these helices break, and . TMS 3 and 8. Binding of the ligand to the outward-facing conformation causes TMS10, the outer gate, to bend and occlude the substrate-binding site. Closure of this gate elicits a transition to the inward-facing conformation by the movement of the hash domain relatively to the bundle domain. The main body movements subsequently lead to the opening of an inner gate and release of Na^+^ and substrate (Simmons et al, 2014; Kazmier et al, 2014; Calabrese et al, 2017). Importantly, structural studies on Mhp1 have been found to be in excellent agreement with functional studies in two homologous subfamilies of NCS1 present in fungi and plants (Krypotou et al, 2012; 2015; Sioupouli et al, 2017; Patching, 2018). In particular, mutations affecting transport kinetics and specificity of members of the so-called Fcy and Fur families have been characterized and shown to be located not only in the substrate binding site, but also in the proposed, by the crystal structures and modeling approaches, outward-facing gate.

We have recently provided genetic evidence that the turnover, function, and intriguingly the specificity of an *Aspergillus nidulans* NCS1 homologue, namely the FurE transporter, depends on interactions of the N- and C-terminal cytoplasmic regions with each other and with the main body of the transporter (Papadaki et al, 2017). In particular, we showed that C- or N-terminally-truncated versions of FurE (FurE-ΔC30 or FurE-ΔΝ21) present increased protein stability under conditions that normally trigger ubiquitylation and endocytic turnover (Vlanti and Diallinas, 2008), but additionally have lost their capacity to import solely one of their three physiological substrates, namely uric acid, while the other two, allantoin and uracil, are transported normally. By isolating genetic suppressors of FurE-ΔC30 restoring uric acid transport, we obtained evidence that the deleted part of the C-terminus has an apparent allosteric effect on the functioning of the substrate translocation trajectory and gating domains. We finally obtained genetic, but also direct evidence using bifluorescence complementation assays that the C-terminus interacts with the distal part of the N-terminus. Our results have suggested that both C- and N-terminal domains are involved in intramolecular dynamics critical for the fine regulation of the mechanism that controls substrate transport.

Here, we dissect further the function of the N- and C-terminal domains of FurE and identify distinct linear segments that are crucial for endocytic turnover, transport activity, substrate specificity or proper folding. Using genetics and Molecular Dynamics (MD) we provide further evidence that specific residues of the N-terminus interact with residues in internal cytoplasmic loops and the C-terminus in a pH-dependent manner, and via these interactions allosterically control the gating mechanism, and thus substrate specificity. Our results are discussed within the context of how the evolution of extended termini in eukaryotic transporters has provided new molecular paths for the generation of novel functions.

## Results

### The N-terminus of FurE is crucial for endocytic turnover, specificity and ER-exit

We have previously characterized the function of FurE-ΔN21, a truncated version of FurE lacking the first 21 amino acid residues from its N-terminus, tagged with a GFP epitope. FurE-ΔN21 proved insensitive to signals triggering endocytosis, such as the presence of ammonium or excess substrates in the growth medium (Gournas et al, 2010; Karachaliou et al, 2013), and consequently the transporter remains stable in the plasma membrane (Papadaki et al, 2017). Most interestingly, FurE-ΔN21 has lost its transport capacity specifically for uric acid, but retains normal transport of allantoin or uracil, as judged from relevant growth tests (purines can be used as sole N sources in *A. nidulans*, whereas uracil transport is scored by sensitivity to the toxic analogue 5-fluorouracil; Papadaki et al, 2017; see also **Figure 1**). Direct transport assays have further shown that loss of transport capacity is not due to reduction in the affinity for uric acid binding, suggesting that inability for uric acid transport is associated with a problem in the dynamics of translocation of this specific substrate.

**Figure 1.**
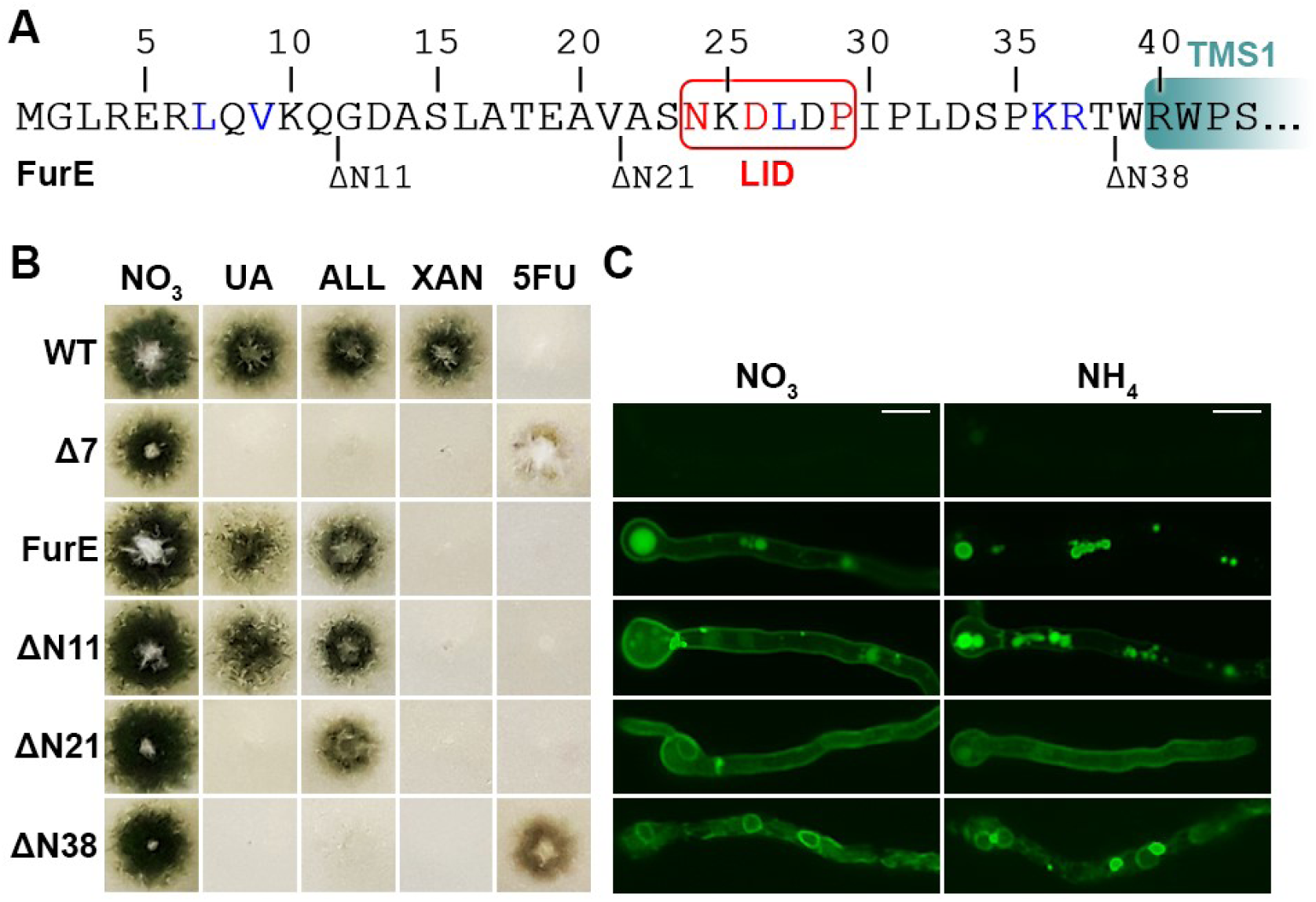
The N-terminus of FurE is crucial for endocytic turnover, specificity and ER-exit. **(A)** Schematic representation of the cytosolic N-terminal region of FurE depicting the limits of deletions ΔN11, ΔN21 and ΔN38, and the conserved LID motif (boxed in red). Conserved amino acids in *Aspergilli* and in all fungal homologues are marked in blue and red respectively. (**B**) Growth test analysis of a standard wild-type (WT) *A. nidulans* strain, a Δ7 strain lacking all genes encoding nucleobase related transporters (*uapAΔ uapCΔ azgAΔ furDΔ furAΔ fcyBΔ cntAΔ*), and isogenic Δ7 transformants expressing functional GFP-tagged ΔN11, ΔN21, ΔN38 and wild-type FurE versions from the strong *gpdA* promoter. The Δ7 strain has an intact endogenous FurE gene transporter, but this is very little expressed under standard conditions and thus does not contribute to detectable transport of its physiological substrates (UA, ALL) or to sensitivity in 5FU (Krypotou et al, 2015). The test was performed on minimal media (MM) containing nitrate (NO_3_), uric acid (UA), allantoin (ALL), xanthine (XAN) as sole nitrogen sources, and NO_3_ plus the toxic nucleobase analogue 5-fluorouracil (5FU), at 37 °C and pH 6.8. (**C**) *In vivo* epifluorescence microscopy of the same strains grown until the stage of young hyphae (16-18 h in MM plus NO_3_^−^). In the right panel ammonium tartrate (NH_4_^+^) was added 2h before microscopic observation. NH_4_^+^-elicited endocytosis is visible as reduced fluorescent signal from the cell periphery concomitant with the appearance of cytosolic structures which correspond to vacuoles and endosomes (Krypotou et al, 2015; Papadaki et al, 2017). For more details see Materials and Methods. Scale bars: 5 μm.

The truncation in FurE-ΔN21 concerned a segment that is little conserved in Fur homologues. However, downstream from this segment a sequence is absolutely conserved in the Fur subfamily, and also well conserved in several prokaryotic NCS1 homologues, including the structurally studied Mhp1. This sequence conforms to the motif: **N**-X-D/S-L-X-**P** (**Figure 1A**). The first Asn and the last Pro are absolutely conserved in all NCS1 members, whereas the Asp-Leu sequence is present in fungal Fur homologues, but replaced by Ser-Asn/Gln in Mhp1 (benzyl-hydantoin transporter) and some other prokaryotic members of various substrate specificities. Given its presence in prokaryotic NCS1 transporters, we predicted that this N-terminal cytosolic motif might be important for the structure and/or function of this group of transporters, rather than being related to membrane traffic, endocytosis or other eukaryotic-specific function. We called this motif, the <u>L</u>oop <u>I</u>nteracting <u>D</u>omain (LID) motif, in accordance with a previous publication referring to it (Keener and Babst, 2013), but also for reasons that will become apparent later.

To investigate the function of the LID segment (residues 21-29) and relate this to the already established role of the distal part of the N-terminus (residues 1-21) in endocytosis and substrate specificity, we constructed and analyzed two new truncated FurE versions. The first was deleted for the entire N-terminus (FurE-ΔN38), and thus for LID too, and the second for the first 11 amino acid residues (FurE-ΔN11). FurE-ΔN38 was found to be retained in the ER membrane and consequently had no apparent transport activity (**Figure 1B, 1C**). In contrast, FurE-ΔN11 possessed normal apparent transport activity and substrate specificity, as judged by growth tests, and showed partial resistance to endocytic internalization, when compared to wild-type FurE and FurE-ΔN21 (**Figure 1B, 1C**). Comparing the effects of the three truncated versions (FurE-ΔN11, FurE-ΔN21 and FurE-ΔN38), it seemed that the distal 11 residues of the N-terminus contribute to endocytosis but are not critical for transport activity or specificity, whereas residues within segment 12-21 also contribute to endocytosis and are crucial for substrate specificity (compare truncations ΔN11 to ΔN21 in respect to endocytosis and growth on uric acid). Finally, segment 22-38 proved critical for ER-exit (compare truncations ΔN21 to ΔN38 in respect to subcellular localization). This analysis however was not sufficient to define the role of the LID as ΔN38 deletion was not sorted to the PM.

### The LID motif determines substrate specificity, but is dispensable for PM localization, transport activity or endocytic turnover

We systemically tested the function of the LID segment, present in the middle part of the N-terminus (residues 21-29), by Ala substitutions of its conserved residues (N24A, D26A, D26A/L27A and P29A). **Figure 2A** shows that LID mutations allow FurE-mediated growth on uric acid and allantoin and confer sensitivity of 5-fluorouracil (5FU), similar to that of an isogenic strain expressing wild-type FurE. Surprisingly, mutations N24A, D26A and especially D26A/L27A conferred growth on xanthine, which normally is not transported by FurE. In other words, Ala substitutions seem not to affect the capacity for transport, but replacements of Asn24 and Asp26/Leu27 enlarged the set of substrates transported to include xanthine. **Figure 2B** confirmed that none of the mutations affects stable localization of FurE into the PM (upper panel), and none has an effect on the sensitivity of FurE to endocytosis elicited by NH_4_^+^ (lower panel). **Figure 2C** showed that the rate of uracil uptake in the mutants is comparable (N24A, P29A) or has a ∼1.5-2-fold increase (D26A, D26A/L27A) compared to that of the wild-type. The accumulation of xanthine by FurE-N24A or FurE-D26A, apparent in growth tests, could not be measured by direct uptake assays, suggesting that this operates by very low-affinity binding of xanthine. Notice that in growth tests purines are added at 400-500 μM, while transport assays are performed with 0.1 μM radiolabeled substrate diluted in non-labeled substrate, so that, very low affinity transport (*K*_m_ > 1mM) cannot be kinetically followed in our system (Krypotou and Diallinas, 2014). The kinetic behavior of N24A, D26A and D26A/L27A mutants, that is, very low affinity transport of a novel substrate, is characteristic of mutations modifying the gating mechanism in other transporters (Papageorgiou et al, 2008; Diallinas, 2016). Thus, our results suggested that the LID sequence affects substrate specificity by altering the mechanism of gating. Notably, while Ala replacements of Asn24 and Asp26 enlarged the set of substrates transported by FurE (allantoin, uracil, uric acid *and* xanthine), deletion of N-terminal residues 1-21 led to restriction of the set substrates transported (uracil and allantoin), as described here and previously (Papadaki et al, 2017). A similar restriction of substrates has also been observed by deleting the last 30 amino acid residues of the C-terminus (Papadaki et al, 2017). These observations will be put in a rational context later.

**Figure 2.**
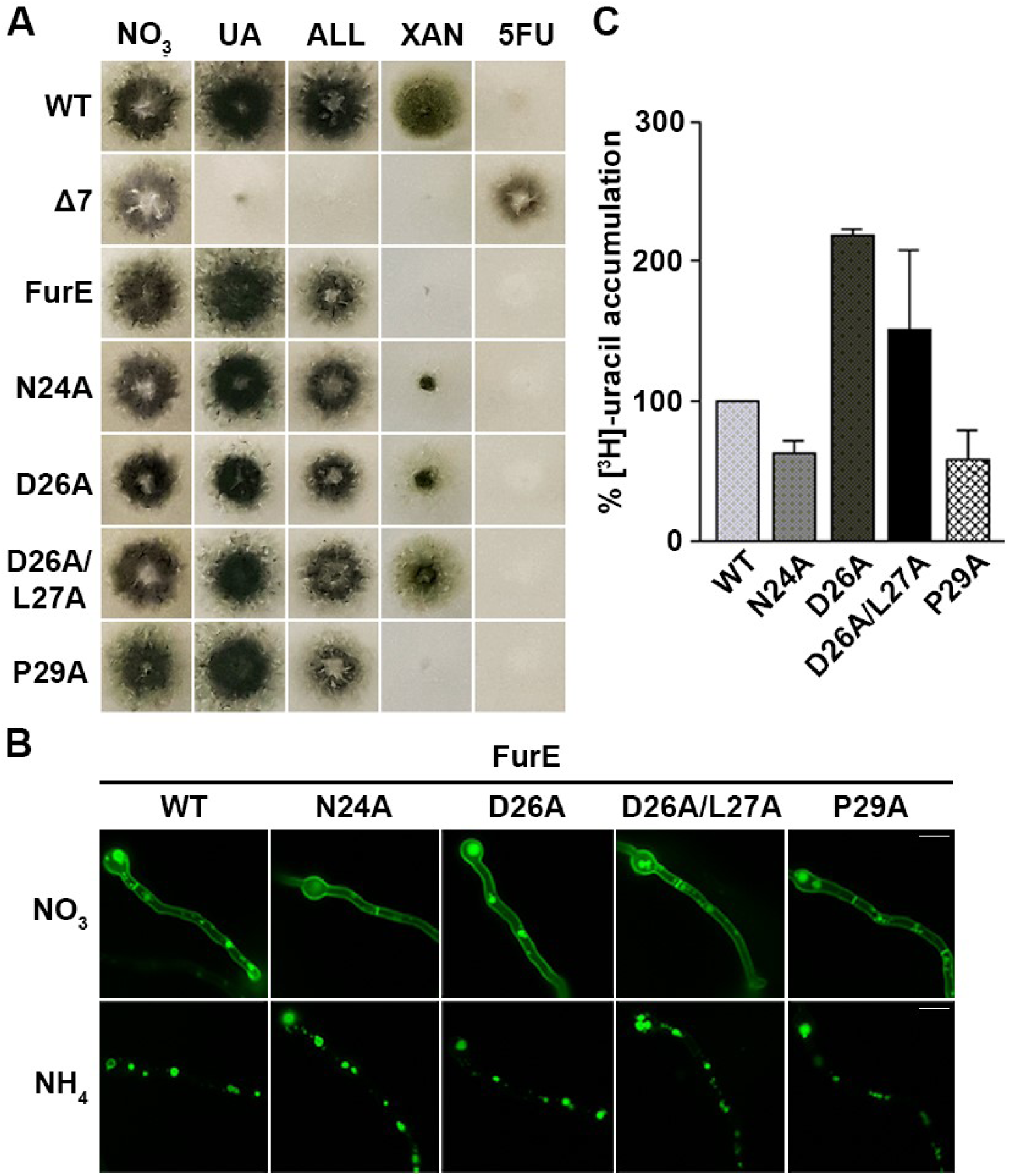
The LID motif of FurE is crucial for substrate specificity, but dispensable for PM localization, transport activity and endocytic turnover. **(A)** Growth tests of the control strains (WT, Δ7, FurE), and strains expressing GFP-tagged FurE mutations in the LID (N24A, D26A, D26A/L27A, P29A). Details are as in Figure 1B. **(B)** Subcellular localization of the FurE mutants. Noticeably, none of the LID mutations affect the PM localization of FurE (NO_3_^−^ panel) or its endocytosis elicited by NH_4_^+^. Details are as in Figure 1C. Scale bars: 5 μm. **(C)** Comparative [^3^H]-uracil accumulation (0.1μΜ) after 2 min of uptake, in strains expressing FurE, FurE-N24A, FurE-D26A, FurE-D26A/L27A or FurE-P29A. Standard deviation is depicted with error bars.

### Delimitation of N-terminal segments crucial for endocytosis, substrate specificity or ER-exit

To obtain a deeper view on the amino acid residues that are crucial for endocytosis versus those that are important for substrate specificity or ER-exit, we systematically mutated, by triple Ala substitutions, the entire N-terminus of FurE. Our results are summarized in **Figure 3**. Growth tests on purines as N sources or on toxic nucleobase analogues revealed that the only triple mutations that led to loss of apparent FurE-mediated transport, as judged by lack of growth on allantoin or uric acid and resistance to 5FU, are those affecting residues 30-32 and 36-38 (**Figure 3A**). This is justified by epifluorescence microscopy, which showed that these two mutations led to retention of FurE in the ER membrane, while all other mutant versions of FurE are properly located in the PM and in some vacuoles, similar to wild-type FurE (**Figure 3B**, see NO_3_ panels). Very minor reduction of growth on allantoin, compared to wild-type FurE, was observed in Ala mutations affecting residues 27-29, but this mutant was still highly sensitive to 5FU, suggesting that its transport activity is generally not affected (**Figure 3A**). Notably, the triple mutations concerning residues 21-23, 24-26, 27-29, and mostly the hexavalent Ala substitution 24-29, conferred growth on xanthine, which is not seen in the control strain lacking FurE (Δ7) or to that expressing a wild-type FurE (**Figure 3A**). This suggests that residues 24-29 of the LID, not only Asn24 and Asp26, are critical for substrate specificity. Additionally, growth on uric acid was reduced, progressively in mutations affecting residues 10-12, 12-14 and mostly 15-17, and also moderately in mutant 33-35 (**Figure 3A**). This means that segments neighboring the LID contribute differentially to substrate selection. We tested whether apparent loss of uric acid uptake in the mutant concerning positions 15-17 (i.e. S15A/L16A, as residue 17 is already an Ala in the wild-type) was due to reduced binding affinity for uric acid. For this, we performed a standard competition assay that measures radiolabel uracil rate in the presence of different concentrations of “cold” uric acid. Our results showed that the *K*_i_ of uric acid for uracil transport competition remains similar to that found for the wild-type FurE (*K*_i_ ∼20 μΜ), suggesting that the defect in the S15A/L16A mutant is uric acid transport *per se*, rather than reduction of the affinity for uric acid binding.

**Figure 3.**
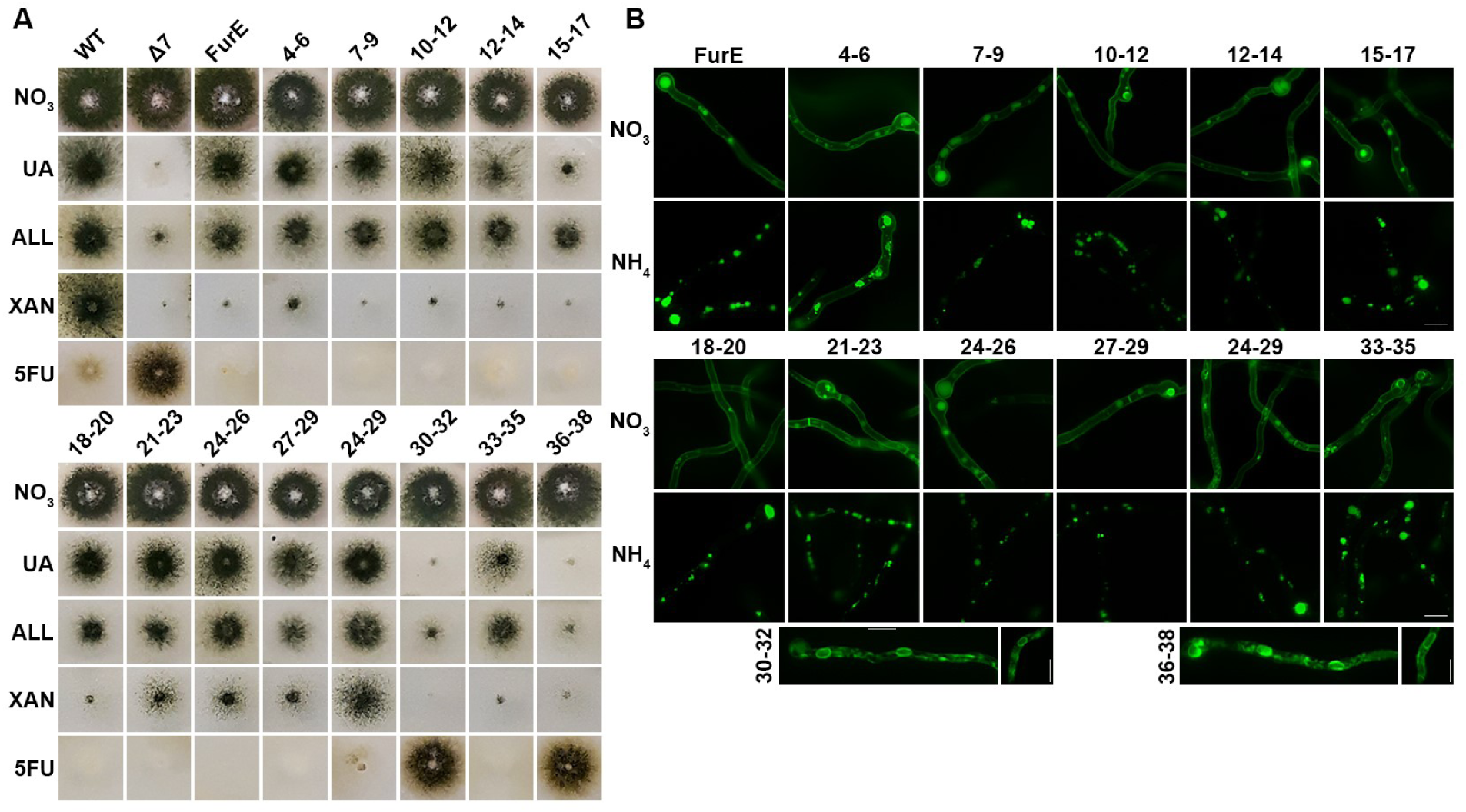
Delimitation of the N-terminal segments crucial for endocytosis, substrate specificity or ER-exit. **(A)** Growth tests of control strains and FurE mutants expressing triple Ala substitutions in the FurE N-terminus. Each mutant is named after the position of residues replaced. Notice that mutant 15-17 has significantly reduced capacity to grow on UA, whereas mutants 21-23, 24-26, 27-29 and mostly 24-29 gained the ability to grow on XAN. Details are as in Figure 1B. (**B**) Epifluorescence microscopy of the mutants shown in A. In the presence of NO_3_^−^ (upper panel) most mutants showed normal PM localization of FurE except mutant 7-9 which showed a degree of instability and increased vacuolar turnover, and mutants 30-32 and 36-38 where FurE was blocked in perinuclear ER rings (Papadaki et al, 2017). Also, all mutants except 4-6, showed the same degree of endocytic turnover as the wild-type FurE after addition of NH_4_^+^. Mutant 4-6 seems to be partially insensitive to endocytosis, marking the PM even in the presence of NH_4_^+^. Details are as in Figure 1C. Scale bars: 5 μm.

In respect to sensitivity to endocytosis, the only Ala triplet that led to detectable but moderately reduced internalization was the one affecting residues 4-6 (**Figure 3B**, lower panel). This contrasts the total block of endocytosis achieved by deleting residues 1-21, and somehow resembles the partial block by deleting residues 1-11 (see **Figure 1**). Noticeably, the distal N-terminal segment of Fur transporter is in general little conserved, but seems to contain an excess of positively charged residue, two of which are removed in in the Ala mutation replacing residues 4-6. Also, given that N- and C-termini have been shown to come into close contact and that their interaction is critical for to endocytosis (Papadaki et al, 2017), the deletion of the N-terminal residues 1-21 might well distruct the interaction with the C-terminus, which in turn affects endocytosis (see also later).

Overall the above mutational analysis showed that N-distal residues (4-6 and 12-21) are essential for endocytosis, middle N-terminal residues affect specificity (mutations in residues 24-29 lead to gain of xanthine transport, whereas mutations in residues 12-17 reduce uric acid transport), whereas residues proximal to TMS1 (30-32 and 36-38) are essential for ER-exit, apparently by affecting proper folding of the transporter.

### Genetic evidence for an allosteric effect of the N-terminus on the functioning of the substrate translocation trajectory

In order to further understand how the N-terminus might affect substrate specificity, we isolated genetic suppressors of mutant S15A/L16A by directly selecting for revertants re-establishing the capacity for FurE-mediated growth on uric acid. Several mutants where selected after U.V. mutagenesis and 24 of them were purified and characterized in respect to their growth phenotypes on purines, and to amino acid changes that occurred within the FurE ORF. **Figure 4A** shows that all suppressor mutations characterized, namely A74V, G291S, S295P, S296R and N308T, confer growth phenotypes on uric acid, allantoin or 5FU similar to that of the isogenic strain expressing wild-type FurE (left panel), but additionally, all except S295P led to weak by clearly detectable growth on xanthine. Furthermore, none of the suppressors affected the proper localization of FurE in the plasma membrane (right panel). All suppressors characterized are located in the TMS7-L7 segment of FurE (**Figure 4B**). and concern changes in well-conserved amino acids. Given that 24 random mutations overall concerned 5 residues, which means that mutagenesis was fairly saturated, and that 20 out of 24 mutations concerned 4 residues located in the same region (TMS7), it seems that the defect caused by the original S15A/L16A mutation (i.e. loss of uric acid transport) is allosterically “transmitted” in the dynamics of TMS7, and this in turn affects substrate translocation in a very specific manner. This region has been suggested to be critical for gating in Mhp1 (Simmons et al, 2014; Kazmier et al, 2014).

**Figure 4.**
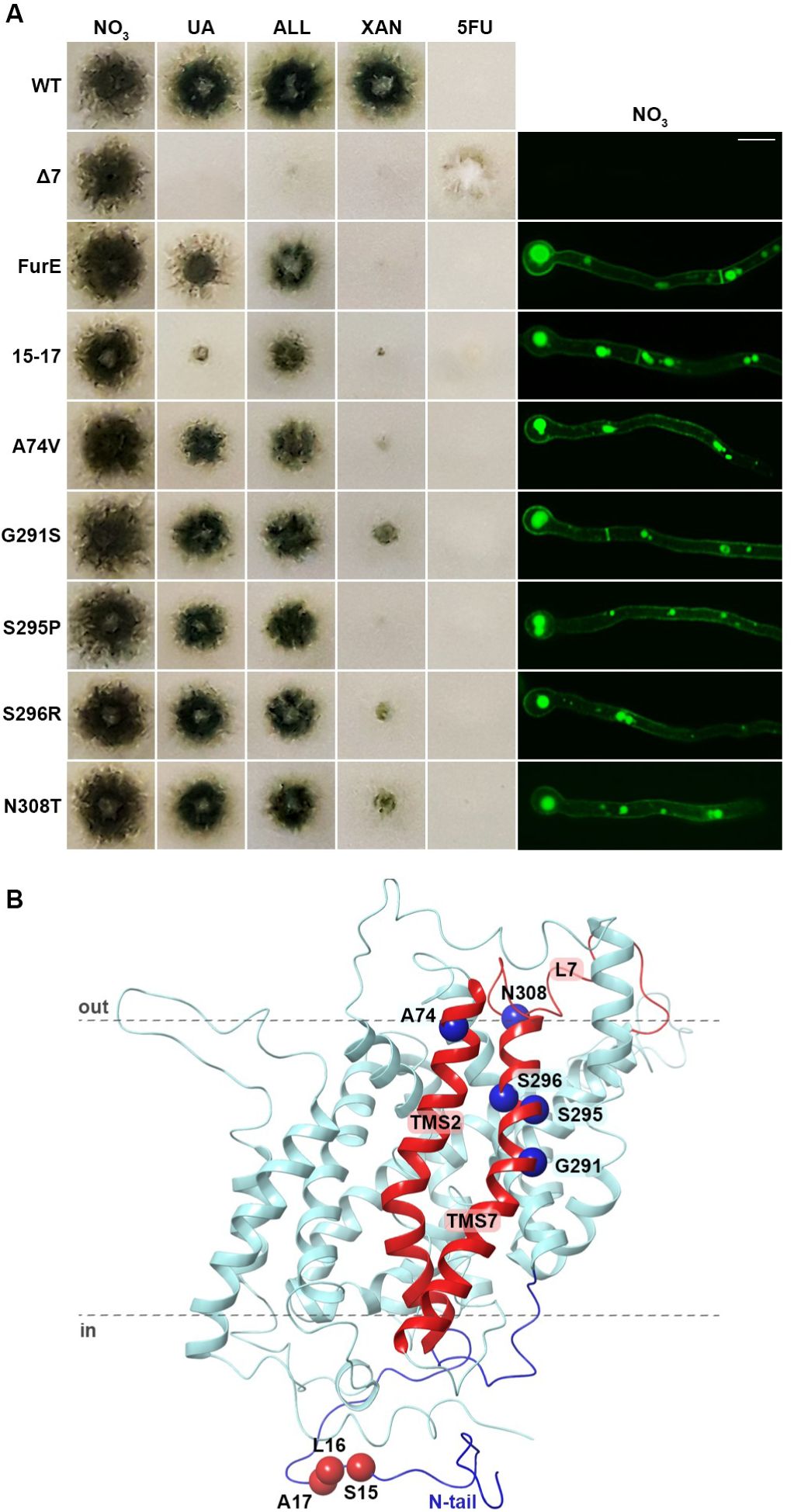
Genetic evidence for an allosteric effect of the N-terminus on the function of the substrate translocation trajectory. **(A)** Growth tests (left panel) and subcellular localization (right panel) of the control strains and FurE-S15A/L16A mutant (named 15-17 in figure) and its suppressor strains. Notice that all suppressors are normally localized in the PM and all have regained the ability of wild-type, FurE-mediated, growth on UA. Details are as in Figure 1B and C. Scale bar: 5 μm. (**B**) Topology of the suppressor mutations (blue spheres) compared to the original N-terminal S15A/L16A mutation (red spheres).

### FurE modeling supports that the N-terminus interacts with specific cytosolic loops and the C-tail

The next question to address was how truncations of terminal segments or amino acid substitutions within the cytosolic N- or C-termini (Papadaki et al, 2017) might be sensed by the transmembrane part of the transporter that hosts the substrate translocation trajectory and carries out transport. Related to this issue, the homologous Mhp1 crystal structure shows that the 20 amino acid region upstream from TMS1, which includes the LID, is in an extended conformation that runs parallel to the membrane along all cytoplasmic loops. Based on this observation, Keener and Babst (2013) have proposed that in *S. cerevisiae* the N-terminus of the homologous Fur4p transporter, and in particular its LID region, might functionally and dynamically interact with several cytoplasmic loops when the transporter acquires an outward-facing conformation. This interaction, being dynamic, might then be disrupted by conformational alteration of the transporter to the inward-facing conformation, elicited by substrate binding. These authors further proposed that dissociation of the LID from the loops renders the N-terminus accessible for Rsp5/Nedd4-type ubiquitination and degradation, which in turn would explain the phenomenon of transport activity-dependent turnover of Fur4p. They extended this idea to propose that the LID is acting as a conformational-sensitive degron that drives turnover under conditions that lead to partial misfolding of Fur4p. Notably, however, their hypothesis was not supported by targeted mutations in the LID motif, which in their case led to no detectable effect on Fur4p stability or function. Interestingly, Razavi et al (2018) very recently showed that the N-terminus of the mammalian dopamine transporter (DAT), which is a structural homologue of NCS1 transporters, also interacts dynamically with specific internal loops and the C-terminus, and thus affects the functioning of DAT. Based on these reports, we tried to obtain evidence as to whether the LID of FurE interacts with the cytosolic loops of the transporter, and thus allosterically affects the dynamics of substrate gating and eventual transport specificity.

To identify possible interactions of the N-terminus with internal loops we built upon homology and validated via Molecular Dynamics (MD) a refined FurE structural model. Mhp1 benzyl-hydantoin permease was used as template in the outward-facing crystal structure (PDB entry 2JLN) according to the alignment presented in **Supplementary Figure 1**. Following the structure of Mhp1 the model represents a 12 α-helix fold with TMS1-10 divided in two symmetric sets oppositely oriented adopting the 5HIRT motif. The overall three-dimensional structure of the FurE model (**Figure 5A**) corresponds to an outward-open conformer, with the transmembrane helices connected with rather short loops except the loop between TMS3 and TMS4 with 21 residues, and the loop separating the core and TMS11–12 is longer (26 residues). The model shows that the side chains of residues critical for transport activity (Krypotou et al., 2015), namely Trp130 (Trp117^Mhp1^), Gln134 (Gln121^Mhp1^) in TMS3, and Asn341 (Asn318^Mhp1^) in TMS8, superimpose exactly with the corresponding in Mhp1, oriented to the substrate-binding cavity of the transporter. The residues in TMS1 Ser54 (Gln42^Mhp1^) and Ser56 (Ala44^Mhp1^) although not conserved, but shown to play a critical role in substrate binding (Krypotou et al, 2015), protrude to the binding cavity, and the same holds true for Lys252, which displays a completely different character from the rest of the NCS1 family members. In most members of the family this residue has aromatic or aliphatic character (Trp220 in Mhp1) and further studies are needed to fully elucidate the function of this variant (Krypotou et al, 2015). Other residues common in all members of the NCS1 family are Trp48 (TMS1), which does not seem to be oriented towards the translocation pathway, and Arg108 in L2, which interacts with His427 in L10 and the backbone carbonyl group of Leu420 in the same loop. Arg108 is positioned in a rather hydrophobic crevice between TMS3 and TMS8 which is closed by the LID residues 22-30. A number of specific and putative dynamic contacts of the residues of the N-terminus, and particularly of the LID region, with residues of internal loops and the C-terminus are summarized in **Supplementary Table 1**, the most important of which are depicted in **Figure 5B**. The main interactions identified are: Asn24 (LID) with Arg108 (L2), Asp26 (LID) with Lys355 (L8), Asp28 (LID) with Lys188 (L4) and Tyr265 (L6). Additionally, Lys188 in L4 seems to interact with L8 (Thr359). Residues 32-39 of the N-terminus may also contact several residues in L2, L6 and L8 and specific residues of the C-tail, such as Met505, and probably Asp506 and Asp507. Overall, the LID residues 24-29 are predicted to interact with L2, L4, L6 and L8, while its downstream region (residues 32-39), which is proximal to TMS1, interacts with L2, L4, L6 and the C-terminus. The C-terminus itself might additionally interact with the FurE core domain, mostly via salt bridges of Glu497 and Glu506 with Arg418 (L10) and Arg270 (L6). These interactions were further validated by more extensive MD (see later). Noticeably, despite the low similarity of FurE and Mhp1, similar interactions between the N-terminus and L2, L6, L8 and L10 and the C-terminus are also observed in Mhp1 (**Supplementary Table 1**). It is interesting to note also that in the case of the outward open conformation of LeuT the N-terminal Arg5^LeuT^ interacts with Asp369^LeuT^ in TMS8 forming a salt bridge, which however is disrupted in the inward conformation, as TMS1a and TMS8 are moving apart (Krishnamurthy and Gouaux, 2012).

**Figure 5.**
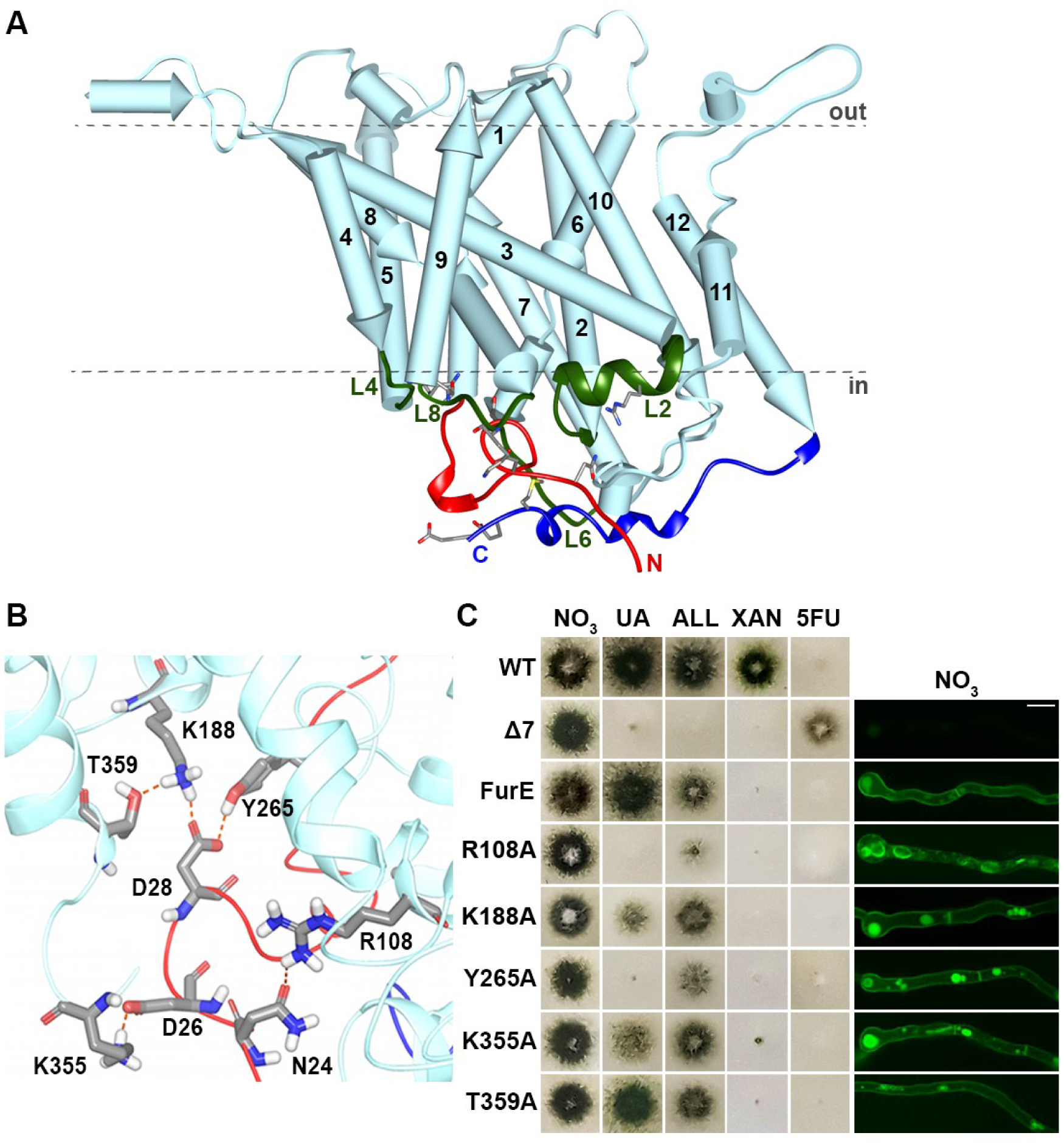
FurE modeling supports that the N-terminus LID might interact with specific cytosolic loops and the C-tail. **(A)** FurE model. Notice the close topological distance of the N-terminal region with internal cytosolic loops L2, L4, L6, L8 and the C-tail. (**B**) Putative major interactions of N-terminal LID with internal cytosolic loops L2, L4 and L8. These interactions were further validated and extended by MD shown in Figure 7 and Supplementary material. **(C)** Growth phenotypes (left panel) and subcellular localization (right panel) of loop mutants R108A (L2), K188A (L4), Y265A (L6), K355A and T359A (L8) and control strains. Details are as in previous figures. Notice that all mutant versions of FurE are localized in the PM, except R108A which is ER-retained. Also, mutations K188A, Y265A and K355A, but not T359A, lead to reduced ability for growth on UA and allantoin, but are still 5FU sensitive, compared to a wild-type FurE. Scale bar: 5 μm.

In case the above proposed interactions are true, relative mutations in specific loop residues might also affect specificity. To experimentally validate this assumption and validate the proposed LID-loops interactions, we constructed and functionally analyzed the following loop mutations; R108A in L2, K188R in L4, Y265A in L6, K355A and T359A in L8. **Figure 5C** shows the growth phenotypes and subcellular localization of the corresponding FurE mutants. Mutation R108A resulted in total ER-retention and thus apparently caused significant FurE misfolding. Thus, no rigrous conclusion on the role of Arg108 in specificity could be drawn. All other mutants were normally localized in the PM and retained at least an apparent normal capacity to transport 5FU. Y265A resulted in reduced allantoin and no uric acid transport, while mutants K188A and K355A had significantly reduced ability to transport uric acid. Finally, T359A showed wild-type transport level for both substrates. In other words, Lys188 (L2), Y265 (L6) and Lys355 (L8) proved crucial for determining the specificity profile of FurE. Noticeably, while mutations within the LID (residues 24-29) enlarged the set of substrates to include xanthine, mutations in loops L2, L6 or L8 restricted the set of substrates to mostly 5FU and allantoin, in a way similar to mutations present just upstream from the LID (residues 12-17). These findings suggest that N-terminus/LID interactions with other internal loops are complex and thus the actual outcome in respect to fine changes in specificity of different Ala substitutions is difficult to predict *a priori*.

### The LID is crucial for determining pH-dependent specificity of FurE

Given that FurE is a proton symporter (Krypotou et al, 2015), we also tested whether its function, and specifically that of the LID motif, might be differentially affected by the proton or cation gradient of the membrane. We tested FurE-mediated growth phenotypes on relevant substrates or toxic analogues in different pHs, as well as, in the presence of a strong Na^+^ gradient. The presence of Na^+^ gradient was found not to affect the apparent function of wild-type or mutant FurE. Contrastingly, we came across a notable pH-dependence of FurE activity, reflected in growth phenotypes, as highlighted in **Figure 6**. In particular, wild-type FurE had significantly reduced apparent transport activity at pH 5.0, as judged by the reduced growth on uric acid and allantoin of the relevant strain, which however could still efficiently transport uracil, reflected sensitivity to 5FU. This contrasts the growth phenotypes observed at pH 6.8, the standard pH where *A. nidulans* is tested (see previous figures). At pH 8.0, FurE confers normal growth on uric acid and allantoin, as well as 5FU sensitivity, but unexpectedly, also leads to significant growth on xanthine, which is not a substrate at pH 6.8 or 5.0 (**Figure 6, middle panel**). Thus, the overall picture is that FurE has a previously unnoticed pH-dependent substrate profile. At low pH it efficiently transports solely 5FU (and apparently uracil), at neutral also pH transports uric acid, and allantoin, while at basic pH additionally transports xanthine.

**Figure 6.**
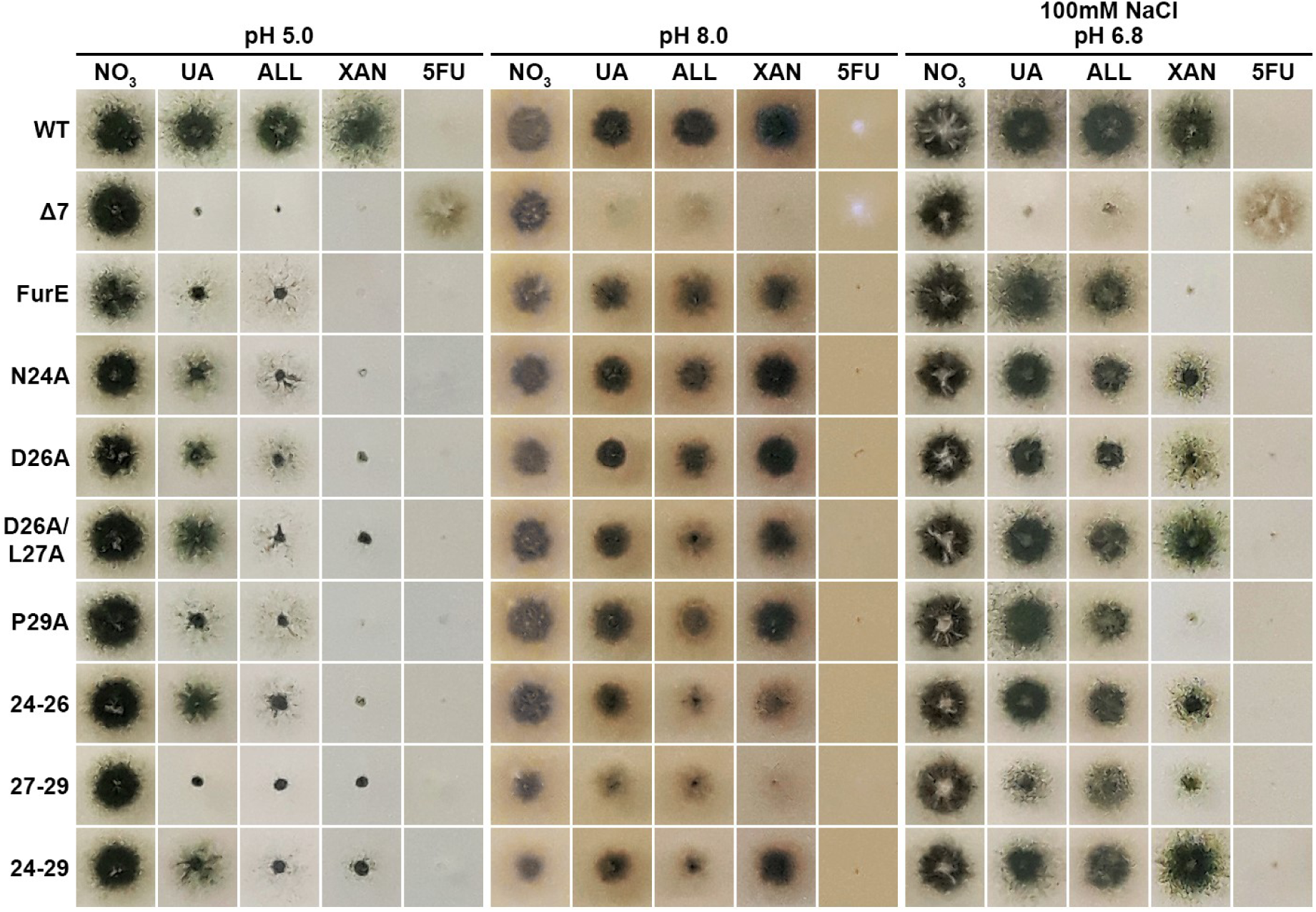
pH dependence of specificity mutations suggest that ion coupling affects LID interactions and gating. Growth tests of control strains and LID mutations in pH 5.0, 8.0 and in pH 6.8 but in the presence of high Na^+^ gradient. For details see Materials and Methods.

Interestingly, the FurE LID mutants also showed distinct pH-dependent phenotypes. At pH 5.0 most LID mutants (N24A, D26A, D26A/L27A, 24-26 and to a lower degree P29A) grow well on uric acid, unlike the strain expressing the wild-type FurE, while their ability to grow on allantoin or uracil remained similar to that of the wild-type FurE. In other words, at low pH LID mutants regain wild-type transport capacity for uric acid, but not for allantoin. At pH 8.0 these mutants conserved the wild-type FurE capacity for xanthine transport, which was also apparent at pH 6.8. These findings revealed that at low pH wild-type FurE functions as a highly specific 5FU (uracil) transporter, incapable for transporting other structurally related substrates, whereas at basic pH FurE becomes an efficient broad-specificity promiscuous transporter, translocating 5FU, uric acid, allantoin and xanthine.

Given that LID mutations showed a distinct pH behavior compared to wild-type FurE, our results suggested that the protonation state of specific residues in the LID, particular the polar or charged residues Asn24, Asp26 or Asp28, might be responsible for the pH-dependent differences in substrate specificity. Most importantly, Ala mutations in the LID somehow mimic the effect of basic pH, leading to increased promiscuity, (i.e. acquisition of the ability to transport xanthine). This observation is discussed further later.

### pH-dependent interactions of the LID with intracellular loops affect gating

In order to gain better insight of the interactions between LID and intracellular facing loops a more detailed structural study has been undertaken by running specific MD calculations that would provide evidence of the flexibility of the LID and the stability of the hydrogen bonds observed in the model. Fundamental to successful MD simulation of a transmembrane protein is the accurate lipid bilayer composition chosen. Here, based on data available for the composition of fungal PM, the lipid bilayer used was composed of 40% phosphatidylcholine 16:1/18:1 (YOPC), 20% Ergosterol (ERG), and 40% phosphatidylinositol lipids (POPI). To specifically address the pH-dependent specificity of FurE, we selected different phosphatidylinositol lipids to emulate different pH environments. For the acidic pH (5.0) POPI lipid models were selected with overall charge −1, (not phosphorylated inositol). For the neutral pH (6.8) 20% POPI and 20% monophosphorylated POPI on position 4 or 5 of inositol with overall charge −3 were mixed (POPI14 or POPI15 equally distributed). Finally, for the basic pH (8.0) we have selected 20% POPI and 20% di-phosphorylated POPI on position 4 and 5 of inositol (POPI24 or POPI25 equally distributed), with overall charge −4. Additionally, we run Molecular Simulations of the FurE mutant version where residues 24-28 of the LID were Ala substituted, using the lipid bilayer simulation for neutral pH (6.8). FurE was embedded on each lipid bilayer, and solvated by explicit water molecules (TIP3P) and 100 ns of simulation have been calculated in all four cases.

The MD simulations reveal that the N-terminus exhibits a dynamic behavior for the part between residues 20-28, that is the LID, while the part proximal to TMS1 (residues 30-40) displays sparse flexibility. The RMSD of the backbone depicted in **Figure 7A** shows that in all cases, except at pH 5.0, the LID is flexible after the first 30 ns period of simulation. **Figure 7B-E** and **Supplementary Video 1** illustrate the significant motion of the LID residues 20-28, and the rather fixed position of downstream residues 29-40 which remain at proximity with mostly L8, but also L2 and L10. It is interesting to notice that although the LID mutant (Ala substituted 24-28) was simulated using the lipid bilayer for neutral pH (6.8) it displays different and more flexible dynamic behavior compared to the wild type, which exhibits only minor deviation from the initial structure for more than the first part of the calculation revisiting positions close to it for the rest of the simulated time. The highest RMSD is attained in the simulation at pH 8.0, reaching 15 Å for most of the calculated time period. This might be due to the higher number of interactions of Lys25 with lipid molecules (**Supplementary Figure 2A**), in addition to a role of other positively charged residues, such as Lys188 and Arg360, which seem to attract PIP2 molecules due to the higher negative charge of the phosphorylated phosphatidylinositol, thus facilitating the displacement of the LID (**Supplementary Figure 2B**). The apparent stability of the segment of residues 29-40 was in good agreement with the experimentally defined structures of Mhp1. The comparison between the two crystal structures, outward-open (2jln) and inward-open (2×79), presented in **Supplementary Figure 3** shows that only the segment upstream from the small bend at Pro15^Mhp1^ is re-oriented in the inward position, thus relaxing the interaction between LID and L8 as TMS8 is also slightly bent, similar to what has been shown for LeuT (Krishnamurthy and Gouaux, 2012). Although there are important differences between Mhp1 and LeuT in the mechanism of substrate translocation it appears that the interruption of the contact between the N-terminus and L8 is common in both cases. Our results suggest that FurE displays the tendency to follow a motion more similar to that observed in Mhp1.

**Figure 7.**
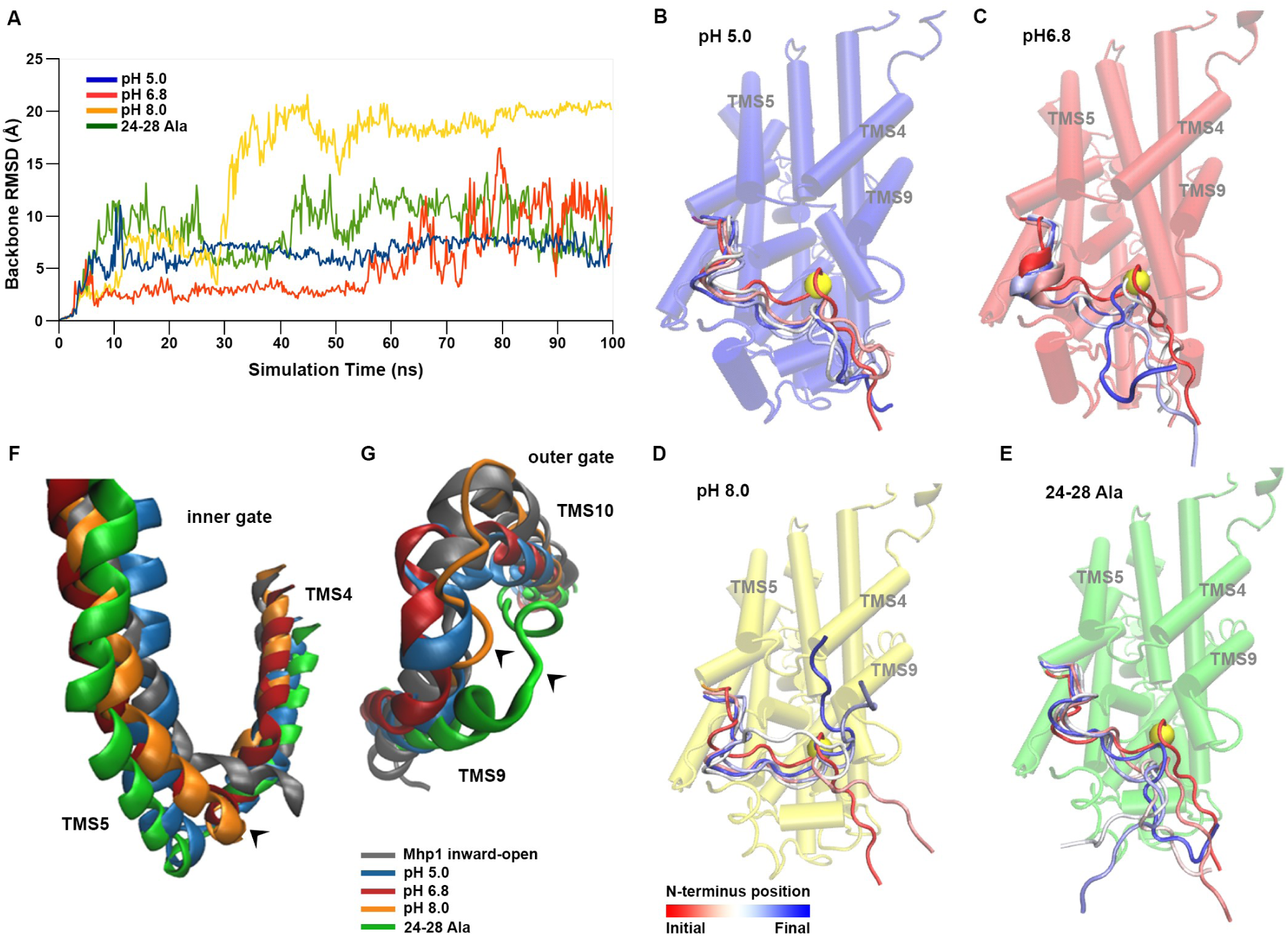
Molecular Dynamics of FurE at different pHs and in the LID mutant. **(A)** RMSD of all Ca atoms of the LID residues 20-40 in respect to the initial structure (blue: pH=5.0, red: pH=6.8, yellow: pH=8 and green: 24-28 Ala mutant). **(B-E)** Schematic representation of FurE cytoplasmic view together with the conformational transition of the LID residues. Snapshots were taken every 25 ns along the transition pathway and are illustrated color code (red for the initial and blue for the final position). Proline 29 showing the residue of the LID most flexible part is labeled as a yellow sphere. **(F-G)** Comparison of the Mhp1 crystal structure (grey) in the inward-open form (PDB 2×79) with the final structures of the four MD simulations of FurE. The two gates are indicated by arrow heads: intracellular gate L4 between TMS4-TMS5 and extracellular gate L9 between TMS9-TMS10. Notice that the outer gate L9 bends covering the binding cavity at pH 5.0 and 6.8, while it remains in open position in the Ala 24-28 mutant. Notice also that the intracellular TMS5 segment shows a propensity to bend, opening the inner gate in the LID mutant, but less apparent in the other three simulated structures.

In order to better visualize and further understand the specific motions of the different helices during the MD simulation we have investigate a) the RMSD of each individual helix, b) the corresponding tilt compared to the Z-axis, and c) the distance of each axis center to that of TMS2, which is the TMS with less motion (**Supplementary Figure 4**). The calculations show that at all pHs and for the LID mutant, helices TMS4, 5, 9 and 10 have a higher propensity to bend, specifically at the loops L4 and L9. The RMSD from the initial position calculated during the MD simulation show that all four TMSs move away from the starting structure between 2 to 5 Å **(Supplementary Figure 4 A-D).** Differences between the four simulations are more pronounced in the case of TMS5, where it seems that in case of simulation of wild-type FurE at pH 8.0 and for the LID mutant (pH 6.8) the deviations are larger than those of wild-type FurE at pH 5.0 and 6.8. Similarly, highest deviations of the initial value are observed for the tilt of the helices compared to the Z-axis are observed in the case of TMS5 with the LID mutant tilting in the opposite direction compared to the three different simulations of pH (**Supplementary Figure 4 E-H)**. Finally, when comparing the distance of the axis between the four TMSs 4, 5, 9 and 10 with TMS2, again the highest variation was observed in the case of TMS5, where the mutant displays the highest deviation from the initial value while the pH 5.0 simulation remains almost stable (**Supplementary Figure 4 I-L)**. Importantly, the specific propensity is clearer when comparing the TMS 5 and TMS9 with the inward open structure of Mhp1. In **Figure 7F-G** the final structures of each one of the four MD calculations are superimposed together with the inward-open Mhp1 structure (2×79). In all four cases, TMS5 shows a propensity to bend towards the inward conformation, with the simulation of pH 8.0 approaching closer to the open structure of Mhp1. In the extracellular interface, TMS9 shows the highest deviation with the LID mutant exhibiting a tendency to remain in the outward-open conformation, while at all pH simulations L9 is bend quite similarly with the Mhp1 inward-open structure.

Overall, MD simulations suggest that the N-terminal LID exhibits relatively high flexibility at the initial part of calculations, more pronounced in the case of pH 8.0 and in the LID mutant, mainly driven from the stronger coulomb interactions between positively charged residues and negatively charged lipids. These interactions mostly influence putative contacts with L8 and TMS9, as shown in **Figure 7F-G.** The proximity of TMS9 to TMS4 appears to be the main reason of a concerted influence to TMS4 and TMS5. The main conclusion from the above MD calculations is that LID motions can influence, in a pH-dependent manner, both the exterior and interior gates and thereby the substrate translocation and transporter specificity. What is also notable is that when the LID motions are higher, as for example is the case at pH 8.0 or in the LID mutant, FurE acts as a promiscuous transporter recognizing all possible substrates, while it becomes more specific for uracil and allantoin at lower pH simulations performed with the wild-type protein.

### Specific C-terminal elements are necessary for endocytosis, transport activity and substrate specificity

In Papadaki et al (2017) we showed that truncation of the 30 last residues of the FurE C-terminus (FurE-Δ498-528) has a dual effect; it blocks endocytosis and leads to a specificity change, in particular loss of uric acid transport. A block of endocytosis of FurE is also achieved when we replace the most distal two lysines in the C-tail (Lys521, Lys522) with Arg resides (**Supplementary Figure 5**). This strongly suggests that block in endocytosis in FurE-ΔC30 is primary due to the lack of these two lysines that apparently act as ubiquitin acceptor residues. To better define the limits of C-terminal segments that affect endocytosis *versus* substrate specificity, we performed systematic Ala replacements of the last 30 residues of the FurE C-tail (**Figure 8A**).

**Figure 8.**
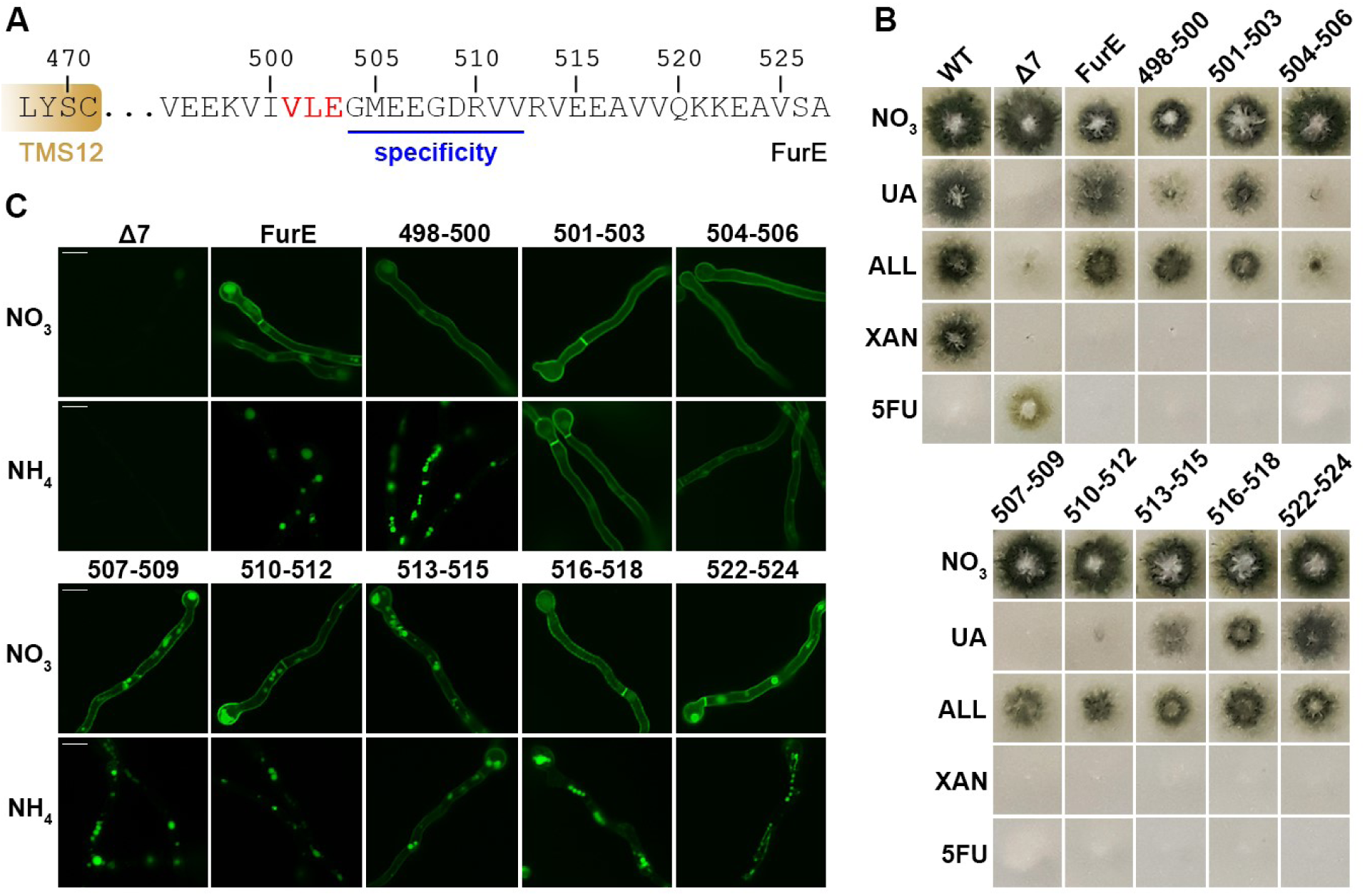
Systematic mutational analysis of the C-terminus defines the limits of elements necessary for endocytosis, transport activity and substrate specificity. **(A)** Sequence of the cytosolic distal C-terminal region of FurE (495-527) The residues involved in endocytosis are highlighted in red and the region involved in substrate specificity determination is underlined, as evidenced in B and C. **(B)** Growth tests of control strains and FurE mutants expressing triple Ala substitutions in the FurE C-terminal region. Each mutant is named after the position of residues replaced. Notice that mutants 504-506, 507-509 and 510-512 have totally lost the ability to grow on UA, while 513-515 has reduced growth on UA. Mutant 504-506 has also significantly reduced ability to grow on ALL and is partially resistant to 5FU, signifying that this mutant is a nearly loss-of-function mutant overall. Details are as in Figure 1B. (**C**) Epifluorescence microscopy of the mutants shown in A. Notice that in the absence of an endocytic signal (NO_3_ panel), all FurE mutants are normally localized in the PM. However, in mutants 501-503 and 504-506, FurE shows increased stability with no sign of steady state vacuolar turnover, as in the wild-type FurE or the other mutants. This is in line with the observation that in these mutants, and particularly in 501-503, FurE is also resistant to NH_4_^+^-elicited endocytosis, suggesting the sequence Val-Leu-Glu is a primary element necessary for endocytosis. Finally, the fact that the following sequence Gly-Met-Glu (504-506) is critical for the transport of all substrates (as shown in B), reveals that this element is absolutely essential for the transport mechanism of FurE *per se*. Scale bars; 5 μm.

**Figures 8B** and **8C** show that triple Ala replacements in the C-terminus affect transport activity, specificity and endocytic turnover of FurE. Ala substitutions of 504-506, 507-509 and 510-512 abolished the ability for growth on uric acid. Mutations in 504-506 also had a strong reducing effect on growth on allantoin and led to partial resistance to 5FU, showing that the sequence Gly-Met-Glu is absolutely essential for transport activity *per se*. Ala substitutions in 513-515 had a moderate effect on uric acid growth. Finally, Ala mutations in the distal C-tail (516-524) had no apparent effect of FurE function. These findings showed that the C-tail is critical for both transport activity (504-506) and specificity 507-512 (**Figure 8A**). In respect to NH_4_^+^-elicited endocytosis, Ala mutations in 501-503 were sufficient to totally block endocytosis, similar to deletion of the C30 segment, while partial abolishment of endocytosis was also observed with Ala mutations in 504-506 and 513-518 (**Figure 8B**). Noticeably, the sequence 501-507 (V-L-E-G-M-E-E) is similar to the C-terminal sequence E/D-X-E-E shown to be essential for ubiquitination and/or endocytosis of the UapA purine transporter (Karachaliou et al, 2013). As the endocytosis of both FurE and UapA is controlled by the same specific α-arrestin adaptor (ArtA), these sequences might be variations of the core target sequence of ArtA.

## Discussion

The importance of cytosolic termini in membrane trafficking processes has been well-documented in several structurally, functionally and evolutionary distinct eukaryotic transporters, such as those specific for amino acids (Crapeau et al, 2014; Ghaddar et al, 2014; Gournas et al, 2016; Popov-Čeleketić et al, 2016), glucose (Kandror and Pilch, 2011; Brodsky 2012; Brewer et al, 2014), carboxylic acids (Becuwe and Leon, 2014; Fujita et al, 2018), L-ascorbic acid (Varma et al, 2009; Kuo et al, 2013), nucleobases (Karachaliou et al, 2013) or nucleosides (Pinilla-Macua et al, 2012; Errasti-Murugarren et al, 2010). In some cases, cytosolic termini are also known to affect the basic transport mechanism. For example, in the Neurotransmitter Sodium Symporter (NSS) family, experimental evidence has suggested that their N- and C-termini are functionally important via their interaction with each other and with other cytosolic loops, as well as, with specific membrane lipids (Guptaroy et al, 2009; Sucic et al, 2010; Cheng and Bahar, 2014; Malinauskaite et al, 2014; Khelashvili, et al, 2015; Sweeney et al, 2017; Razavi et al, 2018). Also, in the case of the yeast Mup1 transporter and the prokaryotic Glu–GABA antiporter GadC, the C-terminus seems to act allosterically on transport activity (Ma et al, 2013; Busto et al, 2018). However, in no case, except FurE (Papadaki et al, 2017), cytosolic termini have been shown to control substrate *specificity*. Here we provide new evidence and a mechanistic rationale on how cytosolic termini affect specificity through fine allosteric regulation of gating. First, using a systematic reverse genetic approach, we topologically and functionally delimit the segments of the N- and C-termini that play distinct roles in trafficking, endocytosis or transport function *per se*. Subsequently, we show, based on unbiased genetic screens and dynamic modeling approaches that N-terminus of FurE interacts with several internal loops and with the C-terminus, and identify specific residues crucial for these interactions by systematic mutational analysis. Finally, we show, through extensive MD simulations that interactions of termini with internal loops are allosterically transmitted to the opening and closing of gates.

Interestingly, N- and C-terminal elements of FurE affected both endocytosis and transport specificity. A critical point in our study was to uncouple the role of distinct terminal elements in these two processes. Thus, endocytosis was found to require, in addition to Lys521 and Lys522 that act as ubiquitylation sites, a short C-terminal sequence Met-Glu-Glu (residues 501-503) and elements within the distal part of the N-terminus (1-21). The need of both terminal segments of FurE for proper endocytic turnover, apparently via a mechanism that involves their dynamic cross-talk (Papadaki et al., 2017), helps to explain how specific conformational changes associated with the transition from an outward to an inward conformation are related with endocytic turnover (Diallinas 2106; Gournas et al, 2016). In a simplified model, when the distal cytosolic termini interact closely with each other and other internal loops, the transporter is found in an outward-facing conformation, while when the interactions of termini and other loops becomes relaxed the transporter is free to alternate in an inward-facing conformation, open to the cytosolic side. This in turn suggests that relaxation of the tight interaction of the N- and C-terminal regions associated with the inward-facing conformation produces a specific conformation more attractive for endocytic turnover. It seems reasonable to propose that N- and C-terminal sequences co-operatively and dynamically regulate the recruitment of the ubiquitylation machinery (e.g. accessibility of arrestin adaptors), which precedes endocytic turnover.

Delimiting the role of terminal elements in FurE endocytosis helped in defining the exact sequences in the two cytosolic termini that are crucial for determining substrate specificity, namely residues 15-31 in the N-terminus and 504-512 in the C-terminus. While work from other groups has shown that cytosolic termini might be critical for overall transport activity, apparently through their effect on the alteration from an outward to an inward topology (Guptaroy et al, 2009; Sucic et al, 2010; Cheng and Bahar, 2014; Malinauskaite et al, 2014; Khelashvili, et al, 2015; Sweeney et al, 2017; Razavi et al, 2018), this does not seem to be the case of FurE, where relevant terminal truncations or mutations affect specificity, rather than overall transport activity. This in turn suggests that specific terminal elements finely regulate the process of gating, rather than the basic alteration from outward to inward conformation. This idea is further supported by the observation that changes in specificity do not seem to be related to significant changes in relevant substrate binding affinities, and mostly by the fact that genetic suppressors of specificity mutations are located along the substrate translocation trajectory in residues potentially acting as gating elements. Our present findings are also in line with previous studies in other transporters, showing that substrate specificity is mostly affected by mutations in gating elements, rather than changes in the major substrate binding site (Diallinas 2016).

How could tails have a distant effect on the transport of a specific substrate, but not of another? The proposed interactions of the N-terminal region, and in particular of the LID, with specific internal loops provide a rationale on how allostery might be transmitted to the opening and closing of gates along the substrate translocating trajectory. More specifically, MD simulations provided evidence that the LID and consequently LID mutations affect, in pH-dependent manner, the relative topology of TMS9-TMS10 (outer gate) and TMS4-TMS5 (inner gate). It should be noted that in our simulations the influence of the charges on the lipid membrane were of crucial importance for approximation of different pHs. Experimental determination of the exact lipid composition of *A. nidulans* PM will be needed to further validate our current approach. Interestingly, FurE proved to function as a rather specific 5FU/uracil transporter at low pH, but as the pH increases FurE becomes progressively more promiscuous, also transporting uric acid and allantoin, and eventually xanthine. In other words, at low pH gating seems stricter, permitting the efficient transport of only 5FU (and apparently uracil), the smallest of FurE substrates, while at higher pHs gating becomes progressively more lose, permitting larger substrates, such allantoin, uric acid and xanthine, to be transported. Given that FurE versions carrying LID mutations mimic the specificity profile of the wild-type FurE at high pH, the relevant mutations seem to lead to loosening of gating, and thus to increased promiscuity in substrate selection. The relative MD analysis of the wild-type and mutant versions of FurE further supported that stricter gating, leading to restricted specificity, is associated with tighter interactions of the cytosolic tails with internal loops, while relaxed gating, leading to transport of increased number of substrates, is associated with loosening of the tight interaction of tails with the main body of the transporter.

## Materials and Methods

### Media, strains and growth conditions

Standard complete (CM) and minimal media (MM) for *A. nidulans* growth were used. Media and supplemented auxotrophies were used at the concentrations given in http://www.fgsc.net. Glucose 1 % (w/v) was used as carbon source. 10 mM ammonium tartrate (NH_4_) or sodium nitrate (NO_3_) was used as nitrogen sources. Nucleobases and analogues were used at the following final concentrations: 5-fluorouracil (5FU) 100 μΜ, uric acid (UA) 0.5 mM, xanthine (XAN) and allantoin (ALL) 1 mM. All media and chemical reagents were obtained from Sigma-Aldrich (Life Science Chemilab SA, Hellas) or AppliChem (Bioline Scientific SA, Hellas). A *ΔfurD::riboB ΔfurA::riboB ΔfcyB::argB ΔazgA ΔuapA ΔuapC::AfpyrG ΔcntA::riboB pabaA1 pantoB100* mutant strain, named Δ7, was the recipient strain in transformations with plasmids carrying *fur* genes or alleles based on complementation of the pantothenic acid auxotrophy *pantoB100* (Krypotou and Diallinas 2014). The Δ7 strain has an intact endogenous FurE gene transporter, but this is very little expressed under standard conditions, and thus does not contribute to detectable transport of its physiological substrates (UA, ALL) or to sensitivity in 5FU (Krypotou et al, 2015). *A. nidulans* protoplast isolation and transformation was performed as previously described (Koukaki et al, 2003). Growth tests were performed at 37 ° C for 48 h, at pH 6.8 or at pH 5.0 and pH 8.0 where indicated.

### Standard molecular biology manipulations and plasmid construction

Genomic DNA extraction from *A. nidulans* was performed as described in FGSC (http://www.fgsc.net). Plasmids, prepared in *E. coli*, and DNA restriction or PCR fragments were purified from agarose 1% gels with the Nucleospin Plasmid Kit or Nucleospin ExtractII kit, according to the manufacturer’s instructions (Macherey-Nagel, Lab Supplies Scientific SA, Hellas). Standard PCR reactions were performed using KAPATaq DNA polymerase (Kapa Biosystems). PCR products used for cloning, sequencing and re-introduction by transformation in *A. nidulans* were amplified by a high fidelity KAPA HiFi HotStart Ready Mix (Kapa Biosystems) polymerase. DNA sequences were determined by VBC-Genomics (Vienna, Austria). Site directed mutagenesis was carried out according to the instructions accompanying the Quik-Change® Site-Directed Mutagenesis Kit (Agilent Technologies, Stratagene). The principal vector used for most *A. nidulans* mutants is a modified pGEM-T-easy vector carrying a version of the *gpdA* promoter, the *trpC* 3’ termination region and the panB selection marker (Krypotou et al,. 2015). Mutations and segment truncations in Fur transporters were constructed by oligonucleotide-directed mutagenesis or appropriate forward and reverse primers (Table S2). Transformants arising from single copy integration events with intact Fur ORFs were identified by Southern and PCR analysis.

### Uptake assays

Kinetic analysis of Fur transporters activity was measured by estimating uptake rates of [^3^H]-uracil uptake (40 Ci mmol^−1^, Moravek Biochemicals, CA, USA), as previously described in Krypotou and Diallinas (2014). In brief, [^3^H]-uracil uptake was assayed in *A. nidulans* conidiospores germinating for 4 h at 37° C, at 140 rpm, in liquid MM, pH 6.8. Initial velocities were measured on 10^7^ conidiospores/100 μL by incubation with concentrations of 0.2–2.0 μΜ of [^3^H]-uracil at 37° C. The time of incubation was defined through time-course experiments and the period of time when each transporter showed linear increased activity was chosen respectively. All transport assays were carried out in triplets. Standard deviation was < 20%. Results were analyzed in GraphPad Prism software.

### Isolation and characterization of suppressor mutations

Suppressor mutations of 10^9^ conidiospores of the strain S15A/L16A were obtained after 3 min 45 sec exposure at a standard distance of 20 cm from an Osram HNS30 UV-B/C lamp and subsequent selection of colonies capable of growing on MM containing uric acid as sole nitrogen source, at 25°C. Spores from positive colonies were collected after 6-8 days and further isolated on the same selective medium that was used to obtain the original colonies. Genomic DNA from 24 purified colonies was isolated and the ORF of FurE was amplified and sequenced. In all cases the amplified fragments contained a new mutation.

### Epifluorescence microscopy

Samples for standard epifluorescence microscopy were prepared as previously described (Gournas et al, 2010; Karachaliou et al, 2013). In brief, sterile 35 mm l-dishes with glass bottom (Ibidi, Germany) containing liquid minimal media supplemented with NaNO_3_ and 0.1% glucose were inoculated from a spore solution and incubated for 18 h at 25°C. The samples were observed on an Axioplan Zeiss phase contrast epifluorescent microscope and the resulting images were acquired with a Zeiss-MRC5 digital camera using the AxioVs40 V4.40.0 software. Image processing and contrast adjustment were made using the ZEN 2012 software while further processing of the TIFF files was made using Adobe Photoshop CS3 software for brightness adjustment, rotation and alignment.

### Homology Modeling

The construction of a structural model of FurE was based on the crystal structure of the Mhp1 benzyl-hydantoin permease from *Microbacterium liquefaciens* in the outward-open structure (PDB entry 2JLN). We utilized as starting alignment the one already described by our group (Krypotou et al, 2015) and optimized based on mutation analysis data (**Supplementary Figure 1**). The final model was built using PRIME software with an energy-based algorithm (Jacobson et al, 2004). A loop refinement routine was also implemented.

### Molecular Dynamics

FurE was inserted into a lipid bilayer using the CHARMM-GUI tool (Wu et al, 2014). The resulting system was explicitly solvated using the TIP3P water model (Jorgensen et al, 1983) and neutralized by the addition of of Na^+^ and Cl^−^ counter ions at concentration of 0.15 M. For the acidic pH (5.0) emulation the lipid bilayer used was composed of 40% phosphatidylcholine 16:1/18:1 (YOPC), 20% Ergosterol (ERG), and 40% POPI lipid models, which were selected with an overall charge −1, (not phosphorylated inositol). For the neutral pH (6.8) emulation the lipid bilayer used was composed of 40% phosphatidylcholine 16:1/18:1 (YOPC), 20% Ergosterol (ERG), and 20% POPI and 20% monophosphorilated POPI on position 4 or 5 of inositol (POPI14 or POPI15 were equally distributed), with an overall charge of −3. Finally, for the basic pH (8.0) emulation, the lipid bilayer used was composed of 40% phosphatidylcholine 16:1/18:1 (YOPC), 20% Ergosterol (ERG), 20% POPI and 20% di-phosphorylated POPI on position 4 and 5 of inositol (POPI24 or POPI25 were equally distributed), with an overall charge of −4. For the FurE LID mutant (24-28 Ala substitution) we utilized the pH 6.8 lipid bilayer emulation described above. Starting from wild type FurE, οn CHARMM-GUI’s initial step “PDB Manipulation Options” we mutated residues 24-28to Alanines. In all cased FurE was embedded on each lipid bilayer and solvated by explicit water molecules (TIP3P). The N-terminal residue (Ala20) was methylated and the C-terminus residue (Glu507) was amidated. All molecular dynamic (MD) simulations were performed with GROMACS 2018 (Abraham et al, 2015) using the all-atom force field CHARMM36 (Huang and MacKerell, 2013). Periodic boundary conditions were used. Long-range electrostatic interactions were treated with Particle Mesh Ewald (PME) method. Non-bonded interactions were described with a Lennard-Jones potential with a cut-off distance of 1 nm and an integration step of 2 fs was implemented. The system was progressively minimized and equilibrated using the GROMACS input scripts generated by CHARMM-GUI and the temperature and pressure was held at 303.15 K and 1 bar respectively (Lee et al, 2016). The resulting equilibrated structures were then used as an initial condition for the production runs of 100 ns with all the constraints turned off. Production runs were subsequently analyzed using GROMACS tools and all images and videos were prepared using VMD software (Humphrey et al, 1996).

## Supporting information

supplementary meterial

## Acknowledgments

This work was supported by a “Stavros S. Niarchos Foundation” grant and by computational time granted from the Greek Research & Technology Network (GRNET) in the National HPC facility -ARIS- under project NCS1_Mechanism (pr006040).

## Author contributions

G.F.P performed all genetic and molecular experiments. A.Z, G.L and E.M performed the *in silico* analysis, the Molecular Dynamics, and analyzed results. E.M. wrote parts of the article. G.D conceived and planned experiments analyzed results and wrote the article.

