## supplementary meterial for "Cytosolic termini of the FurE transporter regulate endocytosis, pH-dependent gating and specificity"

**Supplementary Table 1.** Analysis of putative contacts of the N-terminus of FurE or Mhp1 with the core domain of the transporter. Major interactions (in bold) in FurE are also depicted in Figure 5B.

| FurE Residue | | Electrostatic/Hydrophobic Interactions | Mhp1 Residue | | Electrostatic/Hydrophobic Interactions |
|---|---|---|---|---|---|
| 20 | A | | 8 | E | K398(L10) |
| 21 | V | | 9 | A | |
| 22 | A | | 10 | R | T397(L10) |
| 23 | S | Y421(L10) | 11 | S | Y395(L10) |
| **24** | **N** | **R108(L2)**, L351(TMS8), Y421(L10) | 12 | L | N17(LID),Y395(L10) |
| 25 | K | | 13 | L | R332(L8) |
| **26** | **D** | **K355(L8),** L27, I30(LID) | 14 | N | S16, N17(LID), C327(TMS8), P331(L8) |
| 27 | L | D26(LID) | 15 | P | A18(LID) |
| **28** | **D** | **K188(TMS5), Y265(TMS6)** | 16 | S | N14(LID), F336(TMS9) |
| 29 | P | Y421(L10) | 17 | N | L12, N14(LID),C327(TMS8),Y395(L10) |
| 30 | I | D26(LID) | 18 | A | P15(LID) |
| 31 | P | | 19 | P | |
| 32 | L | R37(LID), G100(L2) | 20 | T | R25(LID), G87(L2), K232(L6) |
| 33 | D | K36, R37(LID) | 21 | R | E24(LID), D464(C-tail), R467(C-tail) |
| **34** | **S** | **M505(C-tail**), R37(LID), R270(L6) | 22 | Y | R25(LID), R85(TMS2), E463(C-tail) |
| 35 | P | | 23 | A | |
| 36 | K | D33(LID) | 24 | E | R21(LID) |
| 37 | R | L32, D33, S34(LID), A97(TMS2), I98(TMS2), A266(L6) | 25 | R | T20, Y22(LID), I84(TMS2), C234(L6) |
| 38 | T | Y265(L6) | 26 | S | E233(L6) |
| 39 | W | V189(L4), Y265(L6), K267(L6), | 27 | V | E233(L6), K235(L6) |

| | | | | | |
|---|---|---|---|---|---|
| | | F262(TMS6) | | | |
| 40 | R | S43, L44, (TMS1) | 28 | G | S31(TMS1), L32(TMS1) |

**Supplementary Table 2.** Oligonucleotides used in this study.

| **Oligonucleotide** | **5'-3' Sequence** |
|---|---|
| GFP NotI R | CGC**GCGGCCGC**TTACTTGTACAGCTCGTCC |
| FurEN11 SpeI F | GCG**ACTAGT**ATGGGCGACGCCAGCCTGGCCAC |
| FurEN21 SpeI F | CGCG**ACTAGT**ATGGCCTCCAACAAAGACCTCG |
| FurEN38 SpeI F | GCG**ACTAGT**ATGTGGAGATGGCCGTCACTAC |
| GFP NotI dstr F | GACGAGCTGTACAAGTAAGCGAACGCGATCCACTTAACGTTACTG |
| GFP NotI dstr R | CAGTAACGTTAAGTGGATCGCGTTCGCTTACTTGTACAGCTCGTC |
| FurE N24A F | GCCACCGAAGCCGTTGCCTCCGCCAAAGACCTCGACCCGATCCCG |
| FurE N24A R | CGGGATCGGGTCGAGGTCTTTGGCGGAGGCAACGGCTTCGGTGGC |
| FurE D26A F | CGAAGCCGTTGCCTCCAACAAAGCCCTCGACCCGATCCCGCTCG |
| FurE D26A R | CGAGCGGGATCGGGTCGAGGGCTTTGTTGGAGGCAACGGCTTCG |
| FurE L27A F | GCCGTTGCCTCCAACAAAGACGCCGACCCGATCCCGCTCGACTCG |
| FurE L27A R | CGAGTCGAGCGGGATCGGGTCGGCGTCTTTGTTGGAGGCAACGGC |
| FurE D26L27A F | CGAAGCCGTTGCCTCCAACAAAGCCGCGGACCCGATCCCGCTCGACTCG |
| FurE D26L27A R | CGAGTCGAGCGGGATCGGGTCCGCGGCTTTGTTGGAGGCAACGGCTTCG |
| FurE P29A F | GCCTCCAACAAAGACCTCGACGCGATCCCGCTCGACTCGCCCAAAC |
| FurE P29A R | GTTTGGGCGAGTCGAGCGGGATCGCGTCGAGGTCTTTGTTGGAGGC |
| FurE DN4-6A F | CACAACTAGTATGGGACTAGCGGCCGCACTCCAAGTAAAACAAGGC |
| FurE DN4-6A R | GCCTTGTTTTACTTGGAGTGCGGCCGCTAGTCCCATACTAGTTGTG |
| FurE DN7-9A F | GTATGGGACTACGAGAAAGAGCGGCCGCAAAACAAGGCGACGCCAG |
| FurE DN7-9A R | CTGGCGTCGCCTTGTTTTGCGGCCGCTCTTTCTCGTAGTCCCATAC |
| FurE DN10-12A F | CGAGAAAGACTCCAAGTAGCGGCCGCCGACGCCAGCCTGGCCAC |
| FurE DN10-12A R | GTGGCCAGGCTGGCGTCGGCGGCCGCTACTTGGAGTCTTTCTCG |
| FurE DN12-14A F | GAAAGACTCCAAGTAAAACAAGCGGCCGCCAGCCTGGCCACCGAAG |
| FurE DN12-14A R | CTTCGGTGGCCAGGCTGGCGGCCGCTTGTTTTACTTGGAGTCTTTC |
| FurE DN15-17A F | GTAAAACAAGGCGACGCCGCGGCCGCCACCGAAGCCGTTGCCTC |
| FurE DN15-17A R | GAGGCAACGGCTTCGGTGGCGGCCGCGGCGTCGCCTTGTTTTAC |
| FurE DN18-20A F | GCGACGCCAGCCTGGCCGCGGCCGCCGTTGCCTCCAACAAAGAC |
| FurE DN18-20A R | GTCTTTGTTGGAGGCAACGGCGGCCGCGGCCAGGCTGGCGTCGC |
| FurE DN21-23A F | GCCTGGCCACCGAAGCCGCGGCCGCCAACAAAGACCTCGACCCG |
| FurE DN21-23A R | CGGGTCGAGGTCTTTGTTGGCGGCCGCGGCTTCGGTGGCCAGGC |
| FurE DN24-26A F | CGAAGCCGTTGCCTCCGCGGCCGCCCTCGACCCGATCCCGCTCG |
| FurE DN24-26A R | CGAGCGGGATCGGGTCGAGGGCGGCCGCGGAGGCAACGGCTTCG |
| FurE DN27-29A F | GTTGCCTCCAACAAAGACGCGGCCGCGATCCCGCTCGACTCGCC |
| FurE DN27-29A R | GGCGAGTCGAGCGGGATCGCGGCCGCGTCTTTGTTGGAGGCAAC |
| FurE DN24-29A F | CCGTTGCCTCCGCGGCAGCCGCAGCCGCGATCCCGCTCGACTCG |
| FurE DN24-29A R | CGAGTCGAGCGGGATCGCGGCTGCGGCTGCCGCGGAGGCAACGG |
| FurE DN30-32A F | CCAACAAAGACCTCGACCCGGCGGCCGCCGACTCGCCCAAACGC |
| FurE DN30-32A R | GCGTTTGGGCGAGTCGGCGGCCGCCGGGTCGAGGTCTTTGTTGG |
| FurE DN33-35A F | CTCGACCCGATCCCGCTCGCGGCCGCCAAACGCACGTGGAGATG |
| FurE DN33-35A R | CATCTCCACGTGCGTTTGGCGGCCGCGAGCGGGATCGGGTCGAG |

| | |
|---|---|
| FurE DN36-38A F | GATCCCGCTCGACTCGCCCGCGGCCGCGTGGAGATGGCCGTCAC |
| FurE DN36-38A R | GTGACGGCCATCTCCACGCGGCCGCGGGCGAGTCGAGCGGGATC |
| FurE R108A F | GCATCAACTTCCCCGTCTACACTGCAGCCAGCTTCGGTATGAAGGG |
| FurE R108A R | CCCTTCATACCGAAGCTGGCTGCAGTGTAGACGGGGAAGTTGATGC |
| FurE K188A F | CAAGCGCCACTGCTATGGCTCGCAGTGTCCAAGCTACGATACC |
| FurE K188A R | GGTATCGTAGCTTGGACACTGCGAGCCATAGCAGTGGCGCTTG |
| FurE Y265A F | CATGCCGGATTTCACGCGGGCCGCCAAAACTCCCAGGGAGGTG |
| FurE Y265A R | CACCTCCCTGGGAGTTTTGGCGGCCCGCGTGAAATCCGGCATG |
| FurE K355R F | CGACCTGGCCCTCTGGTTTCCCAgGTACGTCGATACCCGTCGCG |
| FurE K355R R | CGCGACGGGTATCGACGTACcTGGGAAACCAGAGGGCCAGGTCG |
| FurE T359A F | GGTTTCCCAAGTACGTCGATGCCCGTCGCGGGGCGTATATC |
| FurE T359A R | GATATACGCCCCGCGACGGGCATCGACGTACTTGGGAAACC |
| FurE D498-500A F | CCGTTTGATGTTGAAGAGGCGGCCGCTGTGCTTGAGGGAATGGAGG |
| FurE D498-500A R | CCTCCATTCCCTCAAGCACAGCGGCCGCCTCTTCAACATCAAACGG |
| FurE D501-503A F | GATGTTGAAGAGAAAGTCATTGCGGCCGCGGGAATGGAGGAGGGAG |
| FurE D501-503A R | CTCCCTCCTCCATTCCCGCGGCCGCAATGACTTTCTCTTCAACATC |
| FurE D504-506A F | AAGTCATTGTGCTTGAGGCGGCCGCGGAGGGAGATAGGGTTGTTAG |
| FurE D504-506A R | CTAACAACCCTATCTCCCTCCGCGGCCGCCTCAAGCACAATGACTT |
| FurE D507-509A F | GTGCTTGAGGGAATGGAGGCGGCCGCTAGGGTTGTTAGGGTTGAG |
| FurE D507-509A R | CTCAACCCTAACAACCCTAGCGGCCGCCTCCATTCCCTCAAGCAC |
| FurE D510-512A F | GGAATGGAGGAGGGAGATGCGGCCGCTAGGGTTGAGGAGGCGG |
| FurE D510-512A R | CCGCCTCCTCAACCCTAGCGGCCGCATCTCCCTCCTCCATTCC |
| FurE D513-515A F | GGAGGGAGATAGGGTTGTTGCGGCCGCGGAGGCGGTGGTGCAGAAG |
| FurE D513-515A R | CTTCTGCACCACCGCCTCCGCGGCCGCAACAACCCTATCTCCCTCC |
| FurE D516-518A F | GATAGGGTTGTTAGGGTTGAGGCGGCCGCGGTGCAGAAGAAGGAGG |
| FurE D516-518A R | CCTCCTTCTTCTGCACCGCGGCCGCCTCAACCCTAACAACCCTATC |
| FurE D522-524A F | GGAGGCGGTGGTGCAGAAGGCGGCCGCTGTCTCTGCAGCGGCCG |
| FurE D522-524A R | CGGCCGCTGCAGAGACAGCGGCCGCCTTCTGCACCACCGCCTCC |
| FurE K521/522R F | GTTGAGGAGGCGGTGGTGCAGAGGAGGGAGGCTGTCTCTGCATAG |
| FurE K521/522R R | CTATGCAGAGACAGCCTCCCTCCTCTGCACCACCGCCTCCTCAAC |
| FurE seq 1 | CGCCGTCTTCGGTATGCTTCC |
| FurE seq 2 | CGCGGTACGCCAAAACTCCCAG |
| FurE seq 3 | GCTGCAGTTGGTTGGTGAGC |

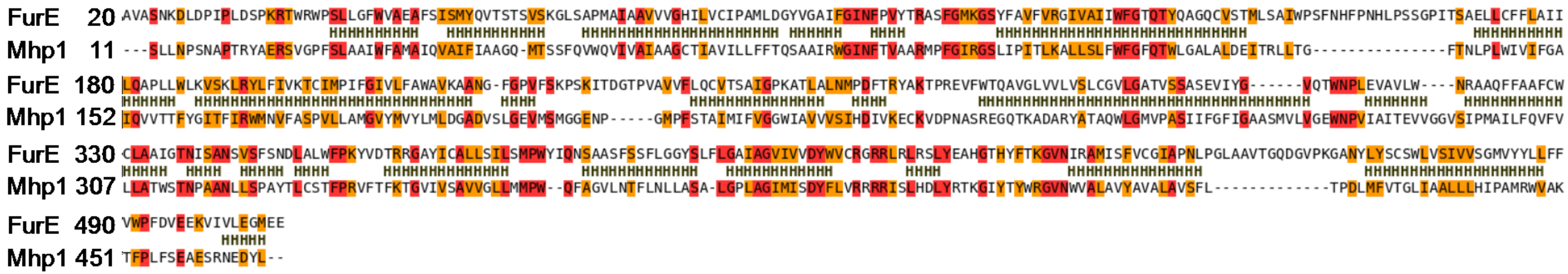

**Supplementary Figure 1.** Binary Alignment of FurE (top) and Mhp1 (bottom) sequence. Helices are depicted with H letter based on 2JLN crystal structure. Color scheme has been applied as implemented on Prime Software for identical (red) or conserved (orange) residues.

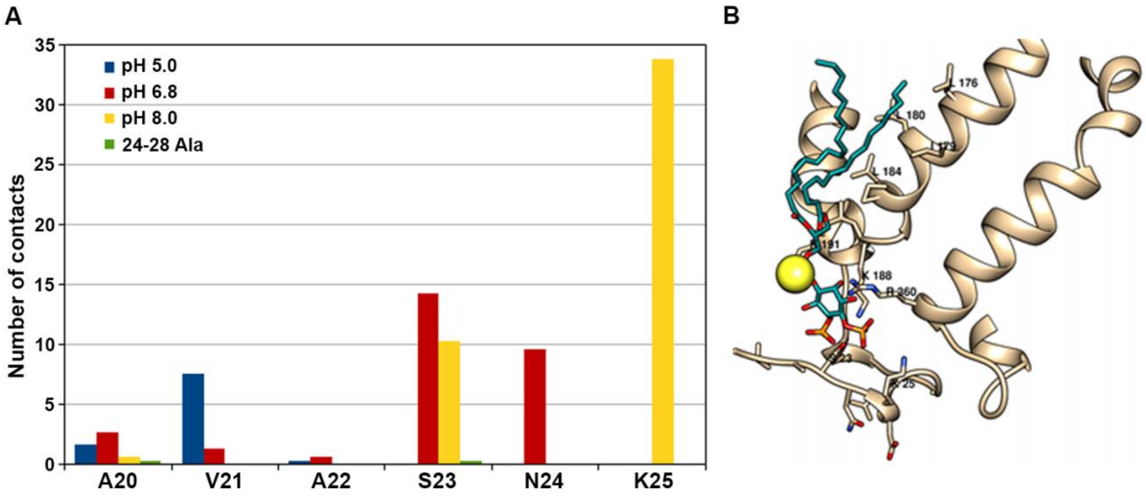


**Supplementary Figure 2**. **(A)** Number of contacts with lipids of different N-terminal residues in all four MD simulations (blue: pH 5.0, red: pH 6.8, yellow: pH 8.0, green: 24-28 Ala mutant). **(B)** Final structure of the MD simulation at pH 8.0 showing the interactions between the 4,5-phosphorylated phosphatidylinositol lipid with FurE. More specifically the polar head forms salt bridges with K188, R360 and K25 and a hydrogen bond with S23. A positively charged sodium ion is also attracted in the vicinity of the above interactions (yellow sphere). The hydrophobic tail of the lipids interacts all along the TMS4 with several lipophilic residue side chains.

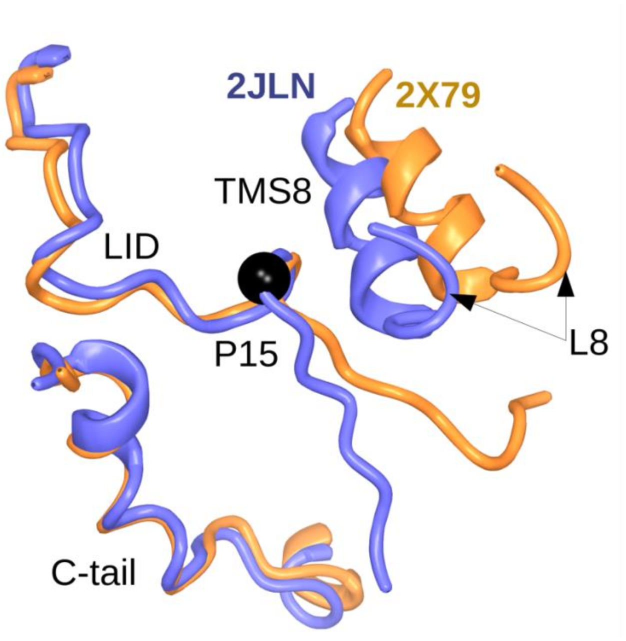


**Supplementary Figure 3.** Superposition of the N-terminus in the outward-open structure (PDB code 2JLN) (blue) onto the inward-open structure (PDB code 2x79) (orange) of the Mhp1 transporter. The L8 loop and TMS8, which participate in interactions with the LID are shown.

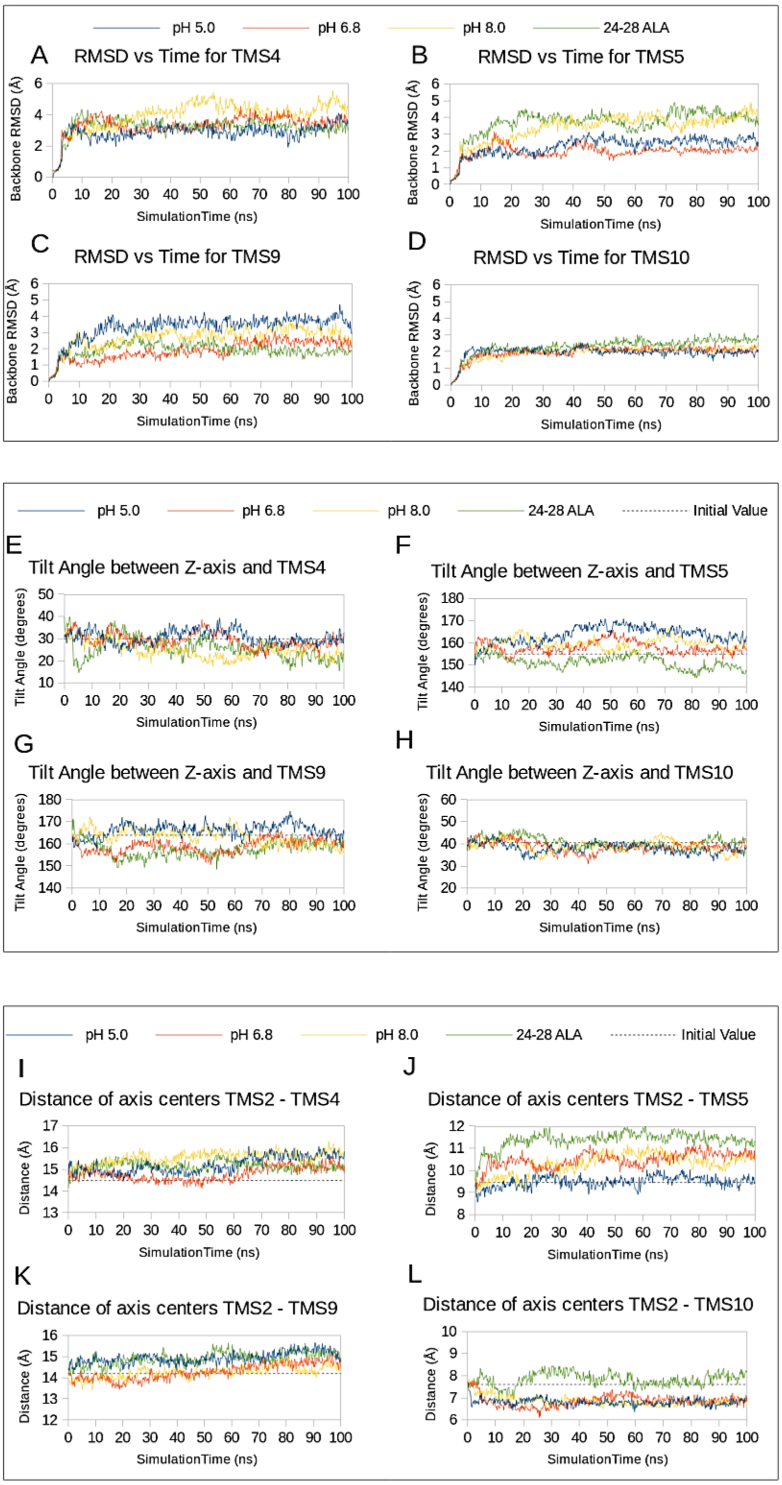


**Supplementary Figure 4.** MD simulation monitoring the motion of TMS4, 5, 9 and 10. (**A-D**) RMSD of all Ca atoms of the specific helix residues in respect to the initial structure versus time. (**E-H**) Tilt Angle between Z-axis and the specific helix. (**I-L**) Distance of axis centers of each specific helix and TMS2. (Blue: pH 5.0, red: pH 6.8, yellow: pH 8.0, green: 24-28 Ala).

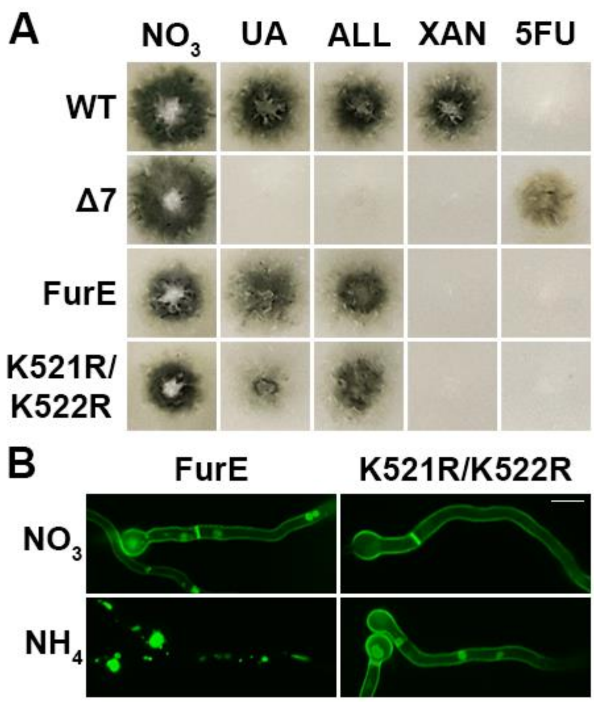


**Supplementary Figure 5. The two more distal Lys residues in the C-terminus of FurE are sufficient for blocking its endocytosis. (A)** Growth test analysis of control strains and FurE K521R/K522R mutant. Details are as in Figure 1B. **(B)** Subcellular localization of the above strains analyzed by in vivo epifluorescence microscopy. FurE genetically lacking the two Lys residues is insensitive to internalization under conditions that trigger endocytosis. Details are as in Figure 1C. Scale bar: 5 μm.

**Supplementary Video legend**

The movie represents 100 ns of Molecular Dynamic Simulation on pH 5.0 (blue), pH 6.8 (red), pH 8.0 (orange) and in the 24-28 Ala LID mutant (green). In all cases the LID is depicted with cyan ribbon. For clarity reasons membrane, solvent molecules and ions are not presented.
